# Extracellular release of mitochondrial DNA is triggered by cigarette smoke and is detected in COPD

**DOI:** 10.1101/2021.10.04.462069

**Authors:** Luca Giordano, Alyssa D. Gregory, Mireia Perez Verdaguer, Sarah A. Ware, Hayley Harvey, Evan DeVallance, Tomasz Brzoska, Prithu Sundd, Yingze Zhang, Frank C. Sciurba, Steven D. Shapiro, Brett A. Kaufman

## Abstract

Chronic obstructive pulmonary disease (COPD) is characterized by continuous and irreversible inflammation frequently caused by persistent exposure to toxic inhalants such as cigarette smoke (CS). CS may trigger mitochondrial DNA (mtDNA) extrusion into the cytosol, extracellular space, or foster its transfer by extracellular vesicles (EVs). The present study aimed to elucidate whether mtDNA is released upon CS exposure and in COPD. We measured cell-free mtDNA (cf-mtDNA) in the plasma of former smokers affected by COPD, in the serum of mice that developed CS-induced emphysema, and in the extracellular milieu of human bronchial epithelial cells exposed to cigarette smoke extract (CSE). Further, we characterized cells exposed to sublethal and lethal doses of CSE by measuring mitochondrial membrane potential and dynamics, superoxide production and oxidative stress, cell cycle progression, and cytokine expression. Patients with COPD and mice that developed emphysema showed increased levels of cf-mtDNA. In cell culture, exposure to a sublethal dose of CSE decreased mitochondrial membrane potential, increased superoxide production and oxidative damage, dysregulated mitochondrial dynamics, and triggered mtDNA release in extracellular vesicles. The release of mtDNA into the extracellular milieu occurred concomitantly with increased expression of DNase III, DNA-sensing receptors (cGAS, NLRP3), proinflammatory cytokines (IL-1β, IL-6, IL-8, IL-18, CXCL2), and markers of senescence (p16, p21). Exposure to a lethal dose of CSE preferentially induced mtDNA and nuclear DNA release in cell debris. Our findings demonstrate that CS-induced stress triggers mtDNA release and is associated with COPD, supporting cf-mtDNA as a novel signaling response to CS exposure.

## Introduction

Chronic obstructive pulmonary disease (COPD) is the third leading cause of death worldwide, excluding COVID-19. It accounts for around three million deaths per year, affects 250 million people worldwide, and is accompanied by social and economic burdens (1). COPD is defined by progressive airflow limitation due to dysregulated chronic inflammation that obstructs the small airways (bronchiolitis) and/or destructs the lung parenchyma (emphysema). Structural remodeling of airway wall thickness and pulmonary vasculature compromise oxygen exchange, leading to long-term disability and early death (2).

The disease is caused by genetic mutations, aberrant cellular responses to bacterial and viral infections, and/or chronic exposure to indoor and outdoor air pollutants, allergens, chemical toxins, and cigarette smoke (CS) – the latter being the most common risk factor (2). Current therapies include bronchodilators, corticosteroids, and antibiotics to slow disease progression, but no existing medications prevent the long-term lung decline. Furthermore, the mechanisms that drive the inflammation and the consequent tissue remodeling in COPD are not completely understood. Notably, a typical feature of chronic inflammation in COPD-affected lungs is the failure to recover even after several years of smoking cessation, suggesting that autoimmunity is a significant driver of the ongoing inflammatory process (3, 4).

Several studies indicate that mitochondrial dysfunction is a hallmark of COPD (5–10). Bioenergetic impairment caused by inhibition of mitochondrial complexes I, III, and IV, overproduction of reactive oxygen species (ROS), and accumulation of dysmorphic mitochondria are common features in cells exposed to CS (5, 6) and in the epithelial cells of lung sections from COPD patients (7, 8). Furthermore, CS-induced lung dysfunction and tissue damage are attenuated by mitochondrial intervention strategies: i) to reduce ROS production by mitochondrial iron chelation or bypassing mitochondrial complex III/IV inhibition (8, 9), or ii) to replace dysfunctional mitochondria by mitochondrial transplantation (10).

Of note, mitochondria are the only extranuclear organelles in animal eukaryotic cells that maintain a genome (mitochondrial DNA, mtDNA). Among its unique features, mtDNA is considered a damage-associated molecule because its mislocalization from the mitochondria to the cytosol or extracellular space (including the blood circulation) can trigger the innate immune system by engaging multiple DNA-sensing receptors (DSRs) and activate downstream proinflammatory signaling cascades. A subset of interactions with major DSRs are described for mtDNA signaling: i) cyclic GMP-AMP synthase (cGAS) recognizes naked mtDNA or mtDNA packaged with mitochondrial transcription factor A (TFAM) (11, 12); ii) Nucleotide-Binding Domain-Like Receptor Protein 3 (NLRP3) binds oxidized mtDNA (13); iii) Toll-Like Receptor 9 (TLR9) recognizes hypomethylated CpG mtDNA sequences (14). Furthermore, Toll-Like Receptor 4 (TLR4) activation during cell injury induces mtDNA fragmentation by endonuclease G, and consequently, contributes to sustaining inflammation (15).

mtDNA damage may be a significant contributor to its mislocalization. mtDNA is a major target of benzo(α)pyrene and its derivatives which are found in CS. (16). In addition, mtDNA is more exposed to oxidative damage because it is more loosely packaged than nuclear DNA (nDNA) and is located near a source of superoxide (17). Alveolar macrophages collected from bronchoalveolar lavage (BAL) from current smokers showed two-fold more nDNA and six-fold more mtDNA damage than non-smokers, as assessed by the percentage of mtDNA common deletions, which is consistent with decreased mtDNA copy number (18). Moreover, a small cohort of lung tissues (n=4) collected from end-stage COPD (GOLD IV) patients showed a substantial increase in mtDNA strand breaks and abasic sites compared to unaffected lungs (19). Because mtDNA damage could be the initiating event in mtDNA release from the mitochondria, these prior reports prompted us to investigate mtDNA mislocalization in models of CS exposure and COPD pathogenesis. Here, we test the hypothesis that CS exposure promotes extracellular mtDNA release, which can be detected in COPD.

## Materials and methods

### Human lung parenchyma and plasma collection

Lung tissues were obtained from 12 COPD patients after lung transplantation at the University of Pittsburgh Medical Center, and normal lung tissues were obtained from 12 donated organs not suitable for transplantation through the Center for Organ Recovery and Education (CORE). Plasma was used from 14 former smokers without lung disease and 20 former smokers diagnosed with COPD randomly selected among the participants in the University of Pittsburgh Specialized Center for Clinically Oriented Research (SCCOR) (20). Demographics and lung physiology are described in **Table E1**. The COPD lung tissue collection (STUDY18100070) and the SCCOR study (STUDY19090239) were approved by the Institutional Review Board for Human Subject Research at the University of Pittsburgh.

### Animal experiments

Animal studies were performed in accordance with the Institutional Animal Care and Use Committee of the University of Pittsburgh (Protocol #18113916). Using a five-chamber smoking apparatus to deliver smoke to mice that were maintained in a monitored, flow-regulated fume hood (21), ten-week-old female mice were exposed to CS from four unfiltered 3R4F reference cigarettes (College of Agriculture, University of Kentucky, Lexington, KY) per day, five days per week, for six months. Control female mice were exposed to room air. Mice were euthanized by CO_2_ inhalation, followed by immediate cardiac puncture to collect the blood, and were then tracheostomized. The right lobe of the lung was dissected and snap-frozen for protein extraction. The left lobe was inflated with 10% formalin at constant pressure for 10 min, fixed, and paraffin-embedded. Blood was allowed to clot in an untreated tube for one h at room temperature and centrifuged at 11,200 × *g* for 10 min at 4°C. Serum (100 μl) was then collected and stored at −80°C.

### Cigarette smoke extract (CSE) preparation

CSE was prepared by burning one 3R4F reference cigarette, bubbling the direct and side-stream smoke into a 50 ml conical tube containing 25 ml of DMEM/F12 (Gibco, Cat#11320-033) without FBS and antibiotics using a peristaltic pump (Precision Blood Pump, COBE Perfusion System) at 0.12 rpm. The CSE-medium was filtered through a 0.22 μm syringe filter (Fisher Scientific, Waltham, MA, USA, Cat#09720004), and the optical density (OD) was measured at a wavelength of 310 nm using a Synergy H1 (BioTek, Winooski, VT, USA) microplate reader. These solutions had OD_310_=0.578 ±0.016, were defined as 100% CSE, then diluted to obtain the desired concentrations (*i.e*., doses). Fresh CSE was prepared for each experiment and used within one h of preparation.

### Statistical analysis

Statistical analysis was performed using GraphPad Prism (version 9.1; GraphPad Software, Inc., San Diego, California). Data are presented as mean±SEM or as median and quartiles (violin or box and whisker plots). After verifying the normality, differences between the two groups were analyzed by Student’s t-test, whereas differences among more than two groups were analyzed by one-way ANOVA with indicated post hoc analyses. Statistical significance was set at *p<0.05, ** p<0.01, ***p<0.001, and ****p<0.0001.

### Additional methods are reported in the online data supplement

## Results

### Cell free-mtDNA and -nDNA are higher in human plasma from former smokers with COPD than former smokers without COPD

Because COPD is characterized by extensive lung inflammation driven by neutrophils, alveolar macrophages, and vascular endothelial cells (4), and mislocalized mtDNA could contribute to inflammation, we determined whether the altered abundance of cell-free DNA could be associated with COPD. We measured cell-free mtDNA (cf-mtDNA) and nuclear DNA (cf-nDNA) in the plasma of former smokers with COPD and former smokers without any clinical evidence of obstruction **(Table E1)**. Both cf-mtDNA and cf-nDNA were elevated in former smokers with COPD compared to the unobstructed former smoker group **(Figure 1A-B)**. In a randomly selected subgroup of the tested plasma, long-range PCR of five overlapping amplicons (encompassing the complete 16.6 kb circular mitochondrial genome) suggested the presence of the intact or large fragments of the mitochondrial genome **(Figure E1A-C)**.

**Figure 1.**
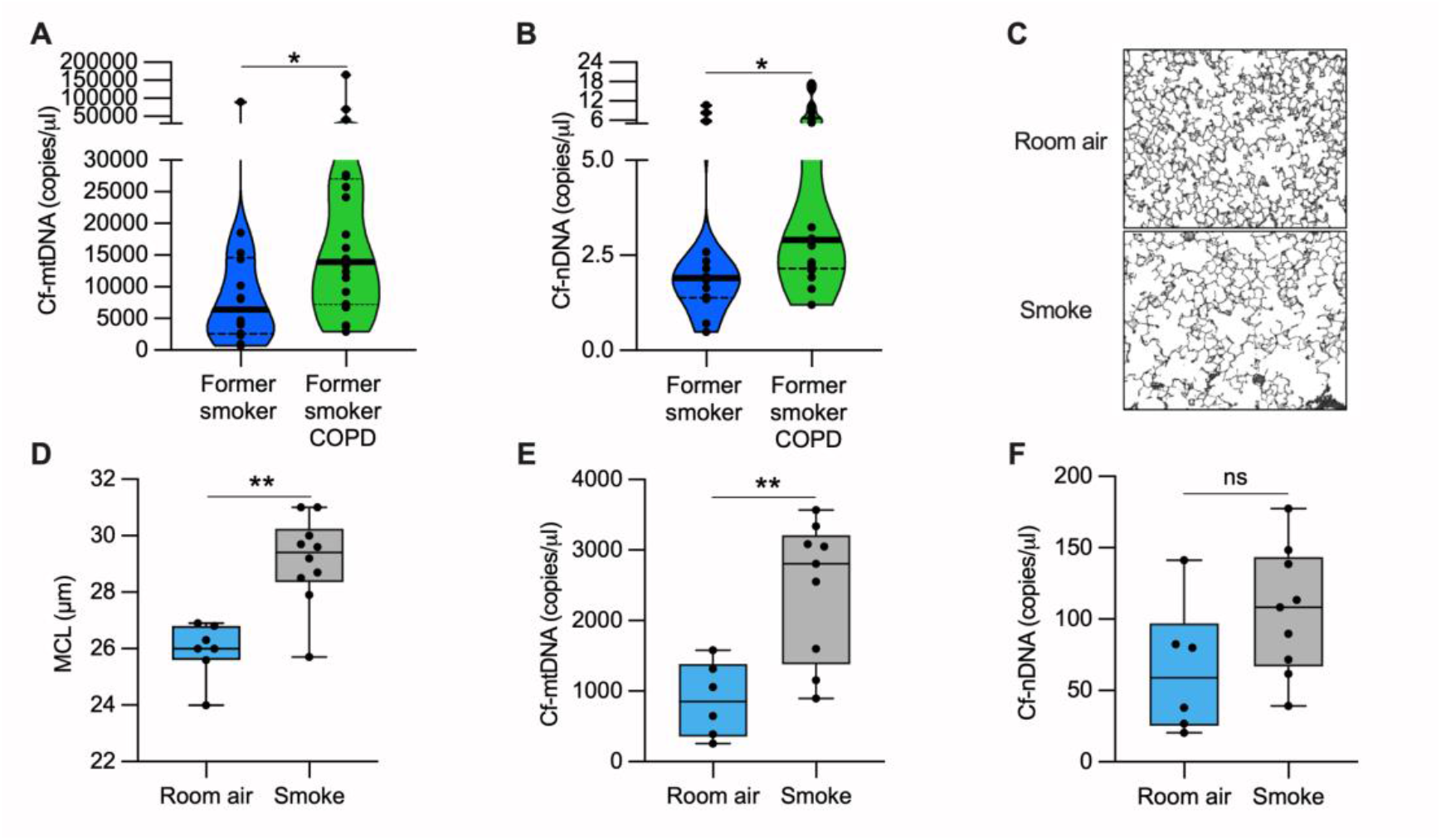
cf-mtDNA is elevated in the plasma of former smokers with COPD and in the serum of a mouse model of emphysema induced by CS exposure. **(A)** Cell-free mitochondrial DNA (cf-mtDNA) and **(B)** cell-free nuclear DNA (cf-nDNA) levels in the plasma of former smokers without airway obstruction (n=14) and former smokers with COPD (n=20). **(C)** Representative images of modified Gill’s stained lung tissue shown in black and white as thresholded for the analysis (original magnification 200X). **(D)** Mean chord length (MCL) quantification of mice exposed to cigarette smoke (n=10) or room air (n=7) for six months. **(E)** cf-mtDNA and **(F)** cf-nDNA in the serum of mice exposed to cigarette smoke (n=9) or room air (n=6) for six months. Data are shown in the (A-B) truncated violin or in the (D-F) box and whiskers plots, with circles representing individual samples. The bold horizontal band in the (A-B) truncated violin plots, represents the median (second quartile), and the dashed bands are the first and the third quartiles. ns, no statistical significance; *p <0.05, **p<0.01 was determined by an unpaired t-test.

Necroptosis has been recently emphasized among several maladaptive pathways involved in COPD pathogenesis (7, 21), including senescence (22), autophagy (23, 24), and apoptosis (24). Necroptosis is a caspase-independent cell death process that promotes a controlled cell membrane lysis and may facilitate DNA release into the extracellular space (25). From lung tissue collected from COPD patients or donors without signs of obstruction, we measured protein levels of the upstream regulators of necroptosis, Receptor-Interacting serine/threonine Protein Kinase 1 and 3 (RIPK1 and RIPK3, respectively). We did not observe a significant change in RIPK1 between the two groups. However, we measured a two-fold increase of RIPK3 in the COPD group compared to the control **(Figure E2A-C)**. Our findings suggest that necroptosis driven by RIPK3 upregulation could be involved in the mtDNA and nDNA release observed in COPD patients.

### cf-mtDNA levels are increased in the serum of mice with alveolar destruction induced by cigarette smoke

To test the cf-mtDNA levels in an animal model, we exposed mice to CS for six months and quantified the lung damage by measuring the mean alveolar chord length. As expected, we found enlarged alveolar airspace (emphysema) in CS-exposed mice compared to the room air-exposed group **(Figure 1C-D)**. In the same cohort of mice, we measured cf-mtDNA and cf-nDNA levels in the serum. CS-exposed mice showed almost three-fold higher cf-mtDNA than the mice exposed to room air **(Figure 1E)**. The cf-nDNA levels followed a similar trend but did not reach statistical significance **(Figure 1F)**. The necroptotic RIPK1 and RIPK3 proteins were higher in the lung lysates of CS-exposed mice than room air controls **(Figure E2D-F)**. These results indicate that, in a mouse model with emphysema, CS exposure upregulates necroptosis markers, and triggers cf-mtDNA extrusion.

**Figure 2.**
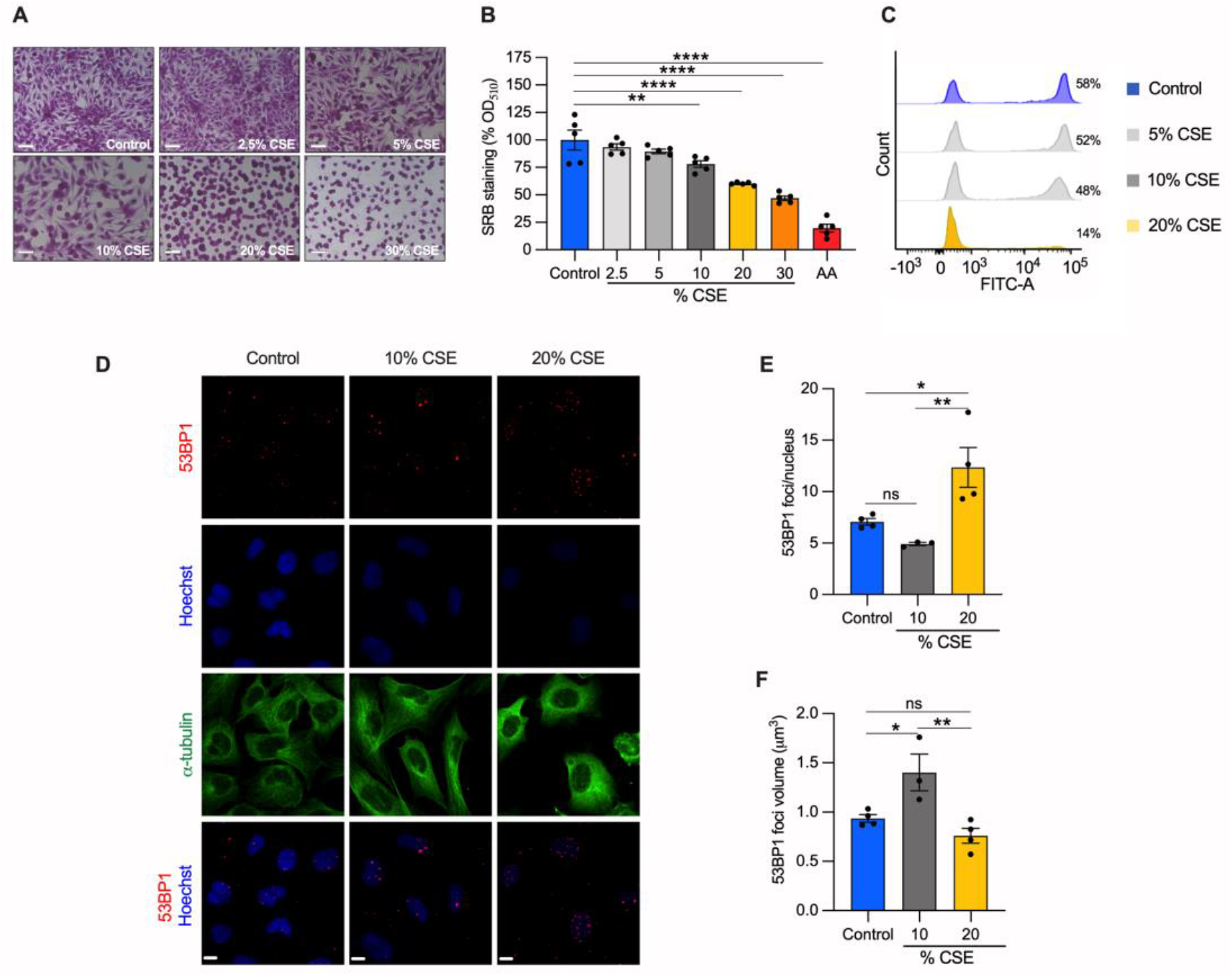
CSE inhibits cell proliferation and promotes cell death through nDNA damage. BEAS-2B cells exposed to indicated CSE dose (%) for 24 hrs were **(A)** stained with Sulforhodamine B (SRB) and **(B)** quantified by measuring the absorbance at 510 nm and normalized to unexposed cells (control, shown as 100%). Antimycin A (AA) was used as a positive control for cell toxicity. Scale bar = 100 μm. **(C)** Representative analysis of flow cytometry showing the number of cells (%) engaged in *de nov*o DNA synthesis by EdU incorporation (n=2). **(D)** Representative maximum projections of confocal image stacks of 53BP1 (red), the nucleus (Hoechst, blue), and the cytoskeleton (α-tubulin, green). **(E)** The number of 53BP1 foci per nucleus, and **(F)** their volume in control, 10%, and 20% CSE-treated cells. Scale bar = 10 μm. (B, E, and F) Data are reported as mean±SEM of independent experiments indicated as circles (n≥3). CSE, cigarette smoke extract; control, CSE-unexposed cells. ns, no statistical significance; *p <0.05, **p <0.01, ****p<0.0001 was determined by one-way ANOVA with multiple comparisons using (B) Dunnett’s or (E-F) Tukey’s post hoc tests.

### Cigarette smoke extract inhibits cell proliferation and promotes cell death in bronchial epithelial cells

To identify molecular mechanisms underlying our *in vivo* observations, we employed an *in vitro* cell model to study CS effects. Specifically, human bronchial epithelial cells (BEAS-2B) were exposed to increasing doses of CSE for 24 hours. Sulforhodamine B assay revealed that CSE toxicity is dose-dependent **(Figure 2A)**. Increased cell death was observed at 20% and 30% CSE, as evidenced by the rounded and shrinking cell morphology. Cells incubated with 10% CSE were spared this severe effect with only a 22% decrease in cell density compared to unexposed cells **(Figure 2A-B)**. To test whether cell death was related to the genotoxic effect of CSE, we measured *de novo* genomic DNA synthesis by 5-ethynyl-2’-deoxyuridine (EdU) incorporation during S-phase. DNA synthesis was almost absent (75.8% decline relative to unexposed) in cells exposed to 20% CSE, whereas cells exposed to 10% CSE showed only a 17.5% decline **(Figure 2C)**. Because DNA double-strand breaks (DSB) induce permanent growth arrest and cell death, we measured the number of p53-binding protein 1 (53BP1) foci, a protein involved in DSB signaling and repair (26). Cells exposed to 20% CSE showed more than double the foci per nucleus than the control **(Figure 2D-E)**. On the contrary, 10% CSE promoted a selective recruitment of 53BP1, as indicated by the increased volume of the foci **(Figure 2D, F)** without increasing foci number. These observations indicate that exposure to 20% CSE mainly triggers cell death (lethal), whereas a lower dose (10%) affects cell proliferation by slowing genomic DNA synthesis (sublethal).

### mtDNA is released into the extracellular milieu via vesicles following exposure to CSE

To test whether DNA is released into the extracellular milieu of bronchial epithelial cells, we exposed cells to increasing doses of CSE for 24 hours, and from the growth medium, we collected extracellular vesicles (EVs) and cell debris. Nanoparticle tracking analysis of the EVs showed no variation in the abundance or size of the particles among the control, 5, and 10% CSE preparations. In contrast, the EVs from 20% CSE preparations were significantly more abundant (1.9×10^9^ *versus* 9.31×10^8^ particles per ml) and slightly larger than the control (150.5 nm *versus* 120.5 nm diameter) **(Figure 3A, E1D)**. In the EVs, mtDNA content increased with the CSE dose, whereas no statistically significant variation of nDNA abundance was detected compared to control **(Figure 3B-C)**. Long-range PCR of the DNA isolated from the EVs showed the presence of the whole or large fragments of the mtDNA genome **(Figure E1E**), in line with the previous results in human plasma **(Figure E1B-C)**. Both mtDNA and nDNA content increased in the cell debris for cells exposed to CSE (5-20%) compared to unexposed cells **(Figure 3E-F)**. Notably, the ratio of mtDNA to nDNA in the EVs increased in a CSE dose-dependent manner **(Figure 3D)** while decreasing in the cell debris **(Figure 3G)**, suggesting distinct mechanisms of DNA release and signaling. These results indicate that exposure to moderate (5-10%) doses of CSE promotes a programmed release of mtDNA by living cells via EVs. On the contrary, both mtDNA and nDNA are effectively released by cell debris due to cell death caused by CSE cytotoxicity.

**Figure 3.**
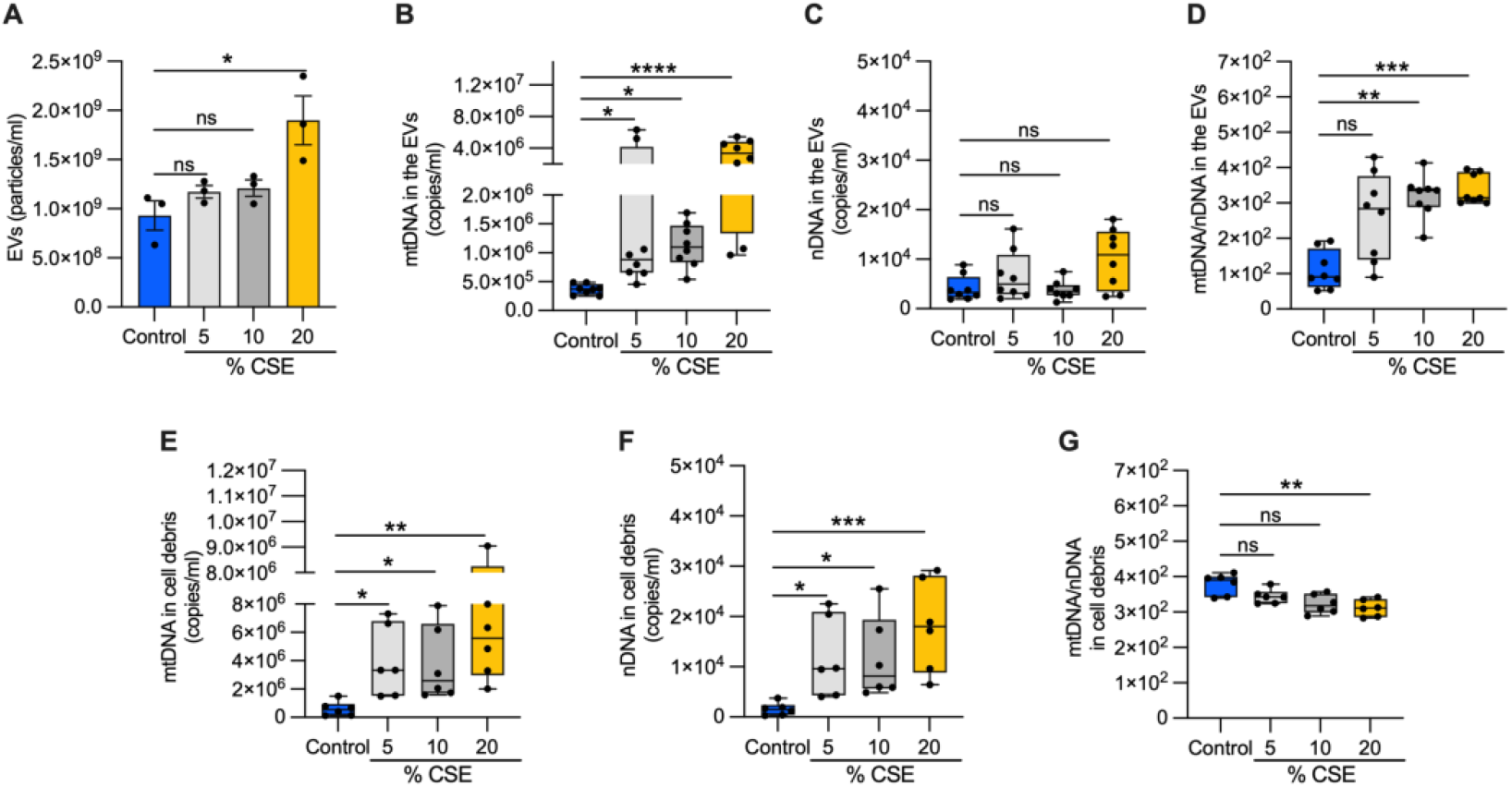
Exposure to CSE promotes mtDNA release in EVs. **(A)** Bar graph showing the EV number (particles/ml) isolated from the extracellular milieu of BEAS-2B exposed for 24 hrs to CSE exposure. Box and whisker plots of the **(B)** mtDNA and **(C)** nDNA content, and **(D)** the mtDNA/nDNA ratio from isolated EVs, as well as the **(E)** mtDNA and **(F)** nDNA content, and **(G)** the mtDNA/nDNA ratio from isolated cell debris. (A) Data are shown as mean±SEM of independent experiments indicated as a circle (n=3). (B-G) In the box and whisker plot, each circle represents the value of an independent experiment (n≥6). CSE, cigarette smoke extract; control, CSE-unexposed cells. (A-G) ns, no statistical significance; *p <0.05, **p <0.01, ***p<0.001, ****p<0.0001 were determined by one-way ANOVA with multiple comparisons and Kruskal-Wallis’s test relative to control.

### CSE promotes mitochondrial membrane depolarization, superoxide production, and oxidative stress

Because mtDNA release could be driven by perturbations of mitochondrial homeostasis (27), we evaluated the effects of CSE on mitochondrial membrane potential and superoxide anion production as direct indicators of mitochondrial dysfunction. We found that CSE promoted mitochondrial membrane depolarization **(Figure 4A)** and increased superoxide production **(Figure 4B)** in a dose-dependent manner. Pretreatment with the mitochondrial scavenger MitoTEMPO prevented superoxide overproduction, confirming that the measurements were specific for the mitochondrial superoxide **(Figure E3A)**.

**Figure 4.**
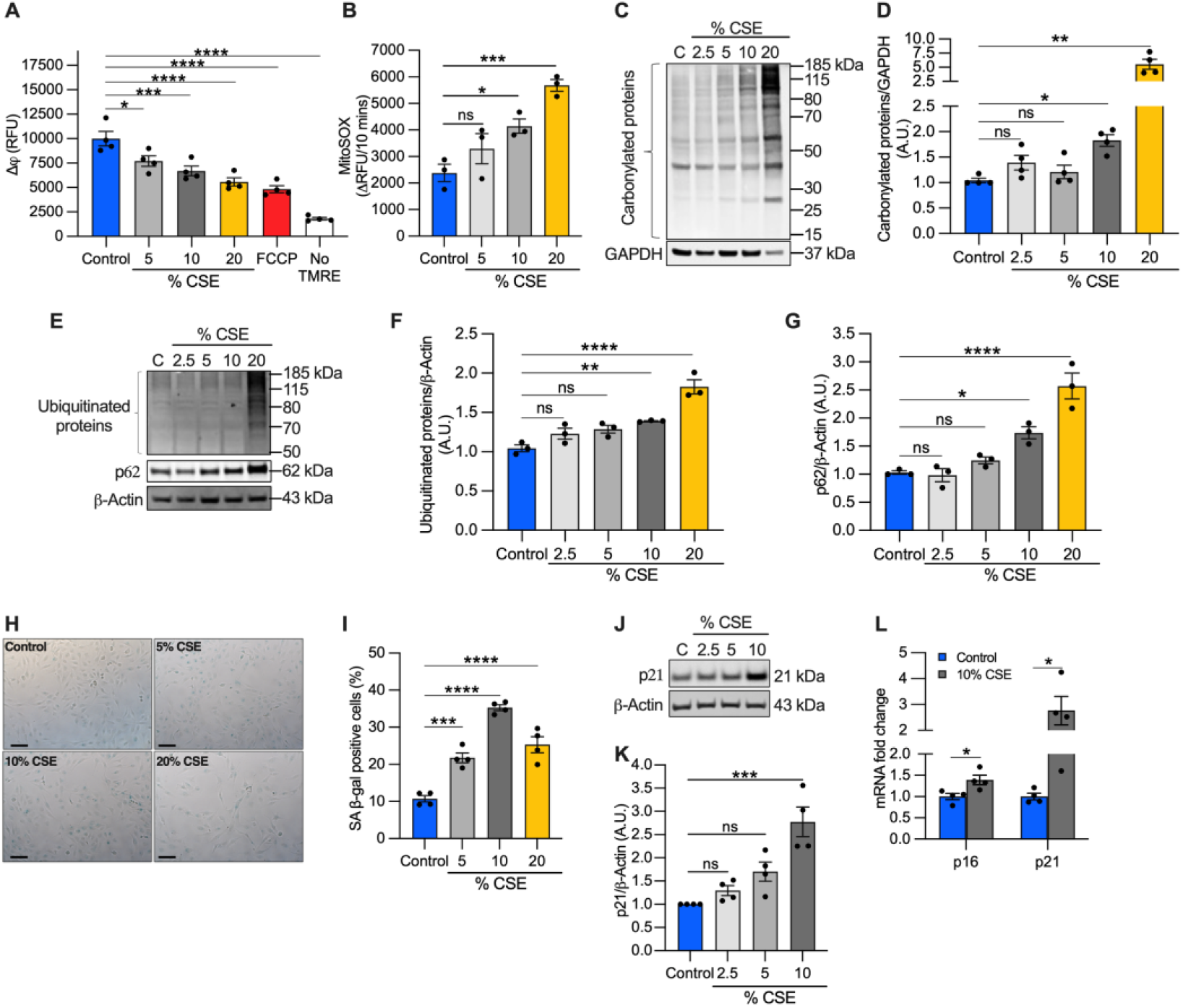
CSE affects mitochondrial membrane potential (Δ*ψ*), promotes oxidative stress, and induces replicative senescence. **(A)** Bar graph reporting the **Δ*ψ*** as relative fluorescence unit of BEAS-2B cells exposed for 6 hrs to CSE and incubated for 20 minutes with 250 nM tetramethylrhodamine ethyl ester (TMRE). Cells incubated without TMRE or with 20 μM carbonyl cyanide p-trifluoro-methoxyphenyl hydrazone (FCCP) were used as negative and positive controls, respectively. **(B)** Bar graph reporting the superoxide anion production as the variation of the relative fluorescence unit (RFU) of MitoSOX over 10 mins in BEAS-2B cells exposed for 3 hrs to CSE. Representative western blots showing total **(C)** carbonylated and **(E)** ubiquitinated proteins, **(E)** p62 and **(J)** p21 proteins in BEAS-2B cells exposed for 24 hrs to CSE. GAPDH or β-actin proteins were used as a loading control. Bar graphs report the densitometry of total **(D)** carbonylated and **(F)** ubiquitinated proteins, and (**G)** p62 and **(K)** p21proteins. **(H)** Representative micrograph images and **(I)** quantification (percentage of total cells) of senescent-associated β-galactosidase positive cells after 24 hrs of exposure to CSE. Scale bar = 100 μm. **(L)** Bar graph reporting the gene expression of p16 and p21 in cells exposed for 24 hrs to sublethal dose (10%) of CSE compared to control. (A-B, D, F-G, I, K-L) Data are shown as mean±SEM of independent experiments (n≥3) indicated as circles. CSE, cigarette smoke extract; control, CSE-unexposed cells. ns, no statistical significance; *p<0.05, **p<0.01, ***p<0.001, ****p<0.0001, (A-B, D, F-G, I) by one-way ANOVA analysis compared to control with Bonferroni post hoc test, or (K) Kruskal-Wallis post hoc tests, and (L) unpaired t-test with Mann-Whitney analysis.

Although moderate levels of superoxide-derived ROS play a role in cellular signaling and adaptation, high ROS levels can lead to protein oxidation and degradation (28). To evaluate the long-term effects of superoxide overproduction, we first quantified carbonylated protein levels, a major product of ROS-mediated oxidation reactions. We found that total carbonylated proteins were modestly increased at 10% CSE, but widely detected at 20% CSE **(Figure 4C-D)**. Next, we measured the total ubiquitinated proteins as a marker of protein degradation, and the levels of the adaptor protein Sequestosome 1 (SQSTM1, *i.e*., p62), which recruits ubiquitinated proteins and organelles to the autophagosome (29). Similar to the results of carbonylation, total ubiquitinated and p62 protein levels increased in cells exposed to 10% CSE and increased further with 20% CSE exposure **(Figure 4E-G)**. These data indicate that CSE exposure affects mitochondrial membrane depolarization, increases superoxide production, and causes protein oxidation and degradation even at sublethal doses (i.e., 10%).

### Exposure to moderate doses of CSE promotes cellular senescence

The mechanisms of mtDNA extrusion remain poorly understood. Specific cell death pathways (necrosis, necroptosis) are proposed to be involved in mtDNA release following severe cell damage (27). Cellular stress induced by CS and CSE has been shown to promote mitophagy-dependent necroptosis (7) and replicative senescence (30). To test whether increased necroptosis was associated with the process of mtDNA release in cells exposed to a sublethal dose of CSE (10%), we investigated the levels of RIPK1, RIPK3, and Mixed Lineage Kinase domain-Like (MLKL, downstream of RIPK1 and RIPK3 in the necroptotic cascade) markers. Both RIPK1 and MLKL proteins showed no substantial changes in abundance in cells exposed to CSE compared to control **(Figure E3B-C, E)**. RIPK3 decreased proportionally to the CSE dose **(Figure E3B, D)** rather than increase as observed in the lung of COPD patients and mice exposed to CS **(Figure E2**).

Because we observed replication inhibition in cells exposed to CSE **(Figure 2C)**, we next tested whether markers of senescence were activated. First, we analyzed cell senescence by β-galactosidase (β-gal) staining. Cells exposed to 5-20% CSE showed an increased number of β-galactosidase positive cells than unexposed cells **(Figure 4H-I)**, with 10% showing peak levels. To confirm replicative senescence in the 10% cells, we measured the proteins levels of the Cyclin-dependent Kinase Inhibitor 1A (p21) as a marker of cell cycle arrest. Western blotting showed a 2.8-fold increase of p21 in cells exposed to 10% CSE compared to control **(Figure 4J-K)**. Both p21 and Cyclin-dependent Kinase Inhibitor 2A (p16) mRNAs were also increased **(Figure 4L)** demonstrating a transcriptional response to block cell cycle progression in cells exposed to 10% CSE. These results suggest that exposure to a sublethal dose of CSE induced mtDNA release in EVs and promotes cellular senescence without activation of necroptosis.

### Mitochondrial dynamics are altered in cells exposed to a sublethal dose of CSE and in COPD lung tissue

Perturbations to mitochondrial dynamics may also favor mtDNA release (27). To test whether the mitochondrial structure was altered, we performed morphometric analysis, using TOMM20 immunostaining to visualize the mitochondrial network. Morphometry showed that the number of mitochondria per cell did not change at 10% CSE **(Figure 5A-B)**, in agreement with the observation that intracellular mtDNA content (mtDNA/nDNA) and TFAM (whose levels usually correlate with mtDNA abundance) were also unchanged **(Figure E3F-H)**. However, the total mitochondrial area and perimeter per cell, and the mean area, perimeter, aspect ratio, and form factor per mitochondrion were increased **(Figure 5A, C-H)**. These results indicate that exposure to a sublethal dose of CSE promotes mitochondrial enlargement and elongation.

**Figure 5.**
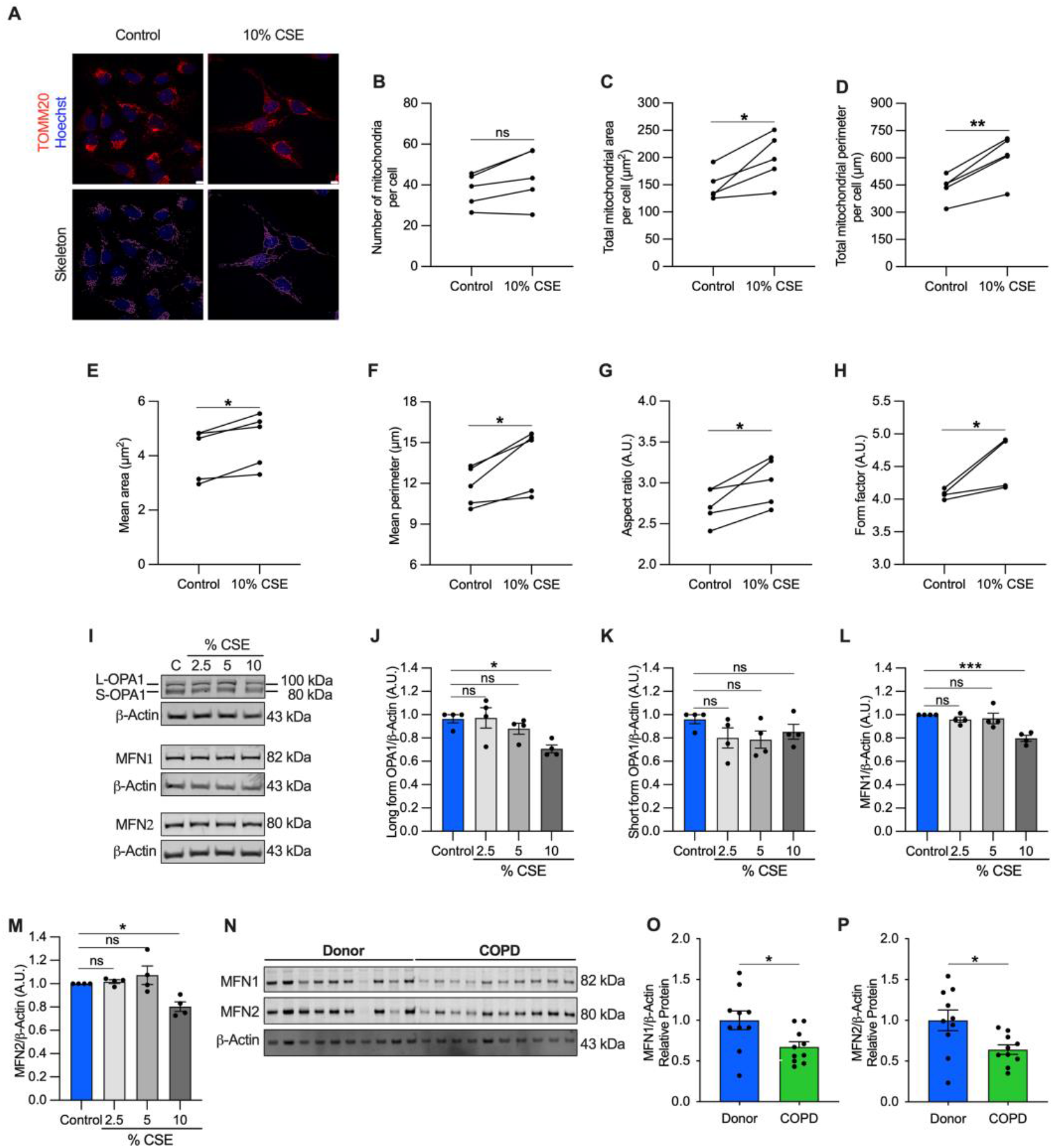
Mitochondrial dynamics is dysregulated in cells exposed to a sublethal dose of CSE and in the airways of the COPD patients. **(A)** Representative maximum projections of 3D-confocal images show the mitochondrial network immunolabeled with TOMM20 (red) in control and cells exposed to 10% CSE for 24 hrs. Nucleus (Hoechst, blue) was used as counterstaining. Greyscale images were converted to binary to generate a skeleton (magenta). Scale bar = 10 μm. Graphs represent **(B)** the number of mitochondria, total mitochondrial **(C)** area and **(D)** perimeter per cell, and the **(E)** mean area, **(F)** mean perimeter, **(G)** aspect ratio (major/minor axis length), and **(H)** form factor (inverse of sphericity) per single mitochondrion. **(I)** Representative western blot of OPA1 (long and short forms), MFN1, and MFN2 protein levels in BEAS-2B cells exposed for 24 hrs to increasing doses of (2.5, 5, 10%) CSE. Bar graphs showing the densitometry of **(J)** long and **(K)** short forms of OPA-1, **(L)** MFN1, and (**M)** MFN2 proteins. **(N)** Representative western blot showing MFN1 and MFN2 proteins, and their relative **(O, P)** densitometry, in the airways of human donors without signs of COPD (n=10) or affected by COPD (n=10). Data are shown as (B-H) mean of 6-8 analyzed images per condition for each independent experiment (n=5), or as mean±SEM of (J-M) independent experiments (n=4), or (O, P) several human samples indicated as circles. CSE, cigarette smoke extract; control, CSE-unexposed cells. ns, no statistical significance; *p<0.05, **p<0.01, ***p<0.001 determined by a (B-H) two-tailed paired t-test, (J-M) one-way ANOVA analysis compared to control with Bonferroni post hoc test, (O, P) unpaired t-test analysis with Welch’s correction.

To enhance our understanding of this process, we quantified protein markers of mitochondrial fusion, including the long-form of Optic Atrophy 1 (L-OPA1), Mitofusin-1 and −2 (MFN1 and MFN2), which are involved in tethering the inner and outer mitochondrial membranes (31). L-OPA1, MFN1, and MFN2 showed a slight but significant decrease in cells exposed to 10% CSE **(Figure 5I-J, L-M)** whereas no changes were observed for the short form of OPA1 (S-OPA1, **Figure 5K)**. Similarly, MFN1 and MFN2 proteins were decreased in COPD lung samples compared to the lung tissue of donors without signs of obstruction, confirming the dysregulated mitochondrial dynamics in the pathological tissues **(Figure 5N-P)**. These findings reveal that alteration of the mitochondrial dynamics is a relevant process in cells exposed to CS and in COPD lung tissue, and could contribute to mtDNA release.

### Exposure to a sublethal dose of CSE upregulates the expression of DNase III, cGAS, NLRP3, and TLR4

The accumulation of mislocalized DNA in our epithelial culture model could be caused by impaired DNases and could lead to the activation of DSRs (32). In cells exposed to 10% CSE, we quantified the expression of the main classes of DNases, differentiated by their activity and localization: DNase I (extracellular), DNase II (phagolysosomal), and DNase III (*i.e*., TREX1; cytosolic) (32). While we did not observe significant differences in the expression of DNases I and II, DNase III showed an almost 50% increase in cells exposed to 10% CSE compared to unexposed cells **(Figure 6A)**, suggesting the activation of a cytosolic mechanism to digest mislocalized DNA.

**Figure 6.**
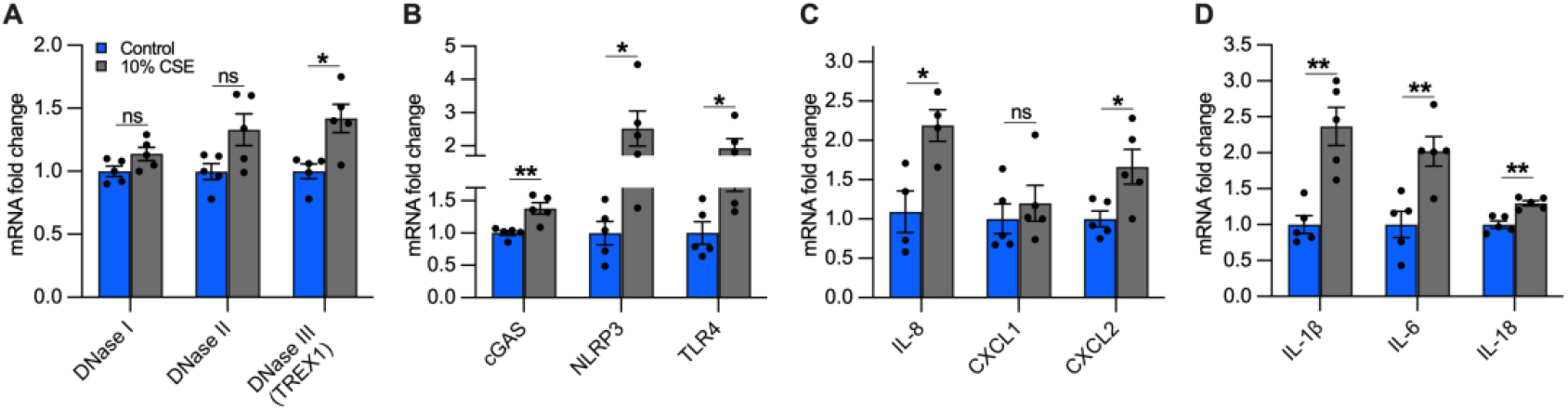
A sublethal dose of CSE upregulates the expression of DNase III, DNA-sensing receptors, and cytokines involved in neutrophil and macrophage recruitment. Bar graphs reporting the gene expression in cells exposed for 24 hrs to a sublethal dose of (10%) CSE compared to control. **(A)** DNases I-III, **(B)** cGAS, NLRP3, TLR4, **(C)** IL-8, CXCL1, CXCL2, and **(D)** IL-1β, IL-6, IL-18. Data are shown as mean±SEM of independent experiments (n≥4), which are indicated as circles. CSE, cigarette smoke extract; control, CSE-unexposed cells. ns, no statistical significance; *p<0.05, **p<0.01 were determined by an unpaired t-test analysis with Welch’s correction.

To identify possible receptors for mislocalized mtDNA, we evaluated the expression of DSRs mainly localized to the cytoplasm that could trigger an inflammatory response caused by CS exposure, specifically cGAS and NLRP3. Furthermore, we measured TLR4, which is involved in mtDNA fragmentation (15), and whose deficiency causes age-dependent emphysema (33). cGAS, NLRP3, and TLR4 expression increased in cells incubated with 10% CSE compared to unexposed cells **(Figure 6B)**. The increased expression of DNase III together with cGAS, NLRP3, and TLR4 suggests that DNA accumulates also in the cytoplasm and may engage DSRs to promote an inflammatory response.

### Exposure to a sublethal dose of CSE upregulates the expression of proinflammatory cytokines involved in the recruitment of neutrophils and macrophages

Inflammation in the lungs of COPD patients is characterized by increased proinflammatory cytokines (4), some of which are downstream products of signaling cascades triggered by mislocalized DNA (11–14). Because neutrophil migration in the airways and inflammation are prominent features of COPD (34, 35), we measured the expression of IL-8, the major chemoattractant of neutrophils in the lungs (35, 36), as well as Chemokine C-X-C motif 1 and 2 (CXCL1 and CXCL2, respectively). Cells exposed to 10% CSE showed an upregulation of IL-8 and CXCL2 expression **(Figure 6C)**, implying a contribution of bronchial epithelial cells to attract neutrophils.

Subsequently, we analyzed the gene expression of IL-1β, IL-6, and IL-18. Chronic expression of IL-1β causes lung inflammation and emphysema by activating lung macrophages (4). IL-6 plays a major role in the systemic inflammation observed in COPD patients, and its overexpression in murine lung cells causes airway inflammation and airspace enlargement. High levels of IL-18 have been found in the lungs of COPD patients and smokers, and its overexpression in mouse lungs triggered inflammatory cell infiltration and alveolar enlargement. In cells exposed to 10% CSE, IL-1β and IL-6 were strongly upregulated while IL-18 was increased to a lesser extent **(Figure 6D)**. Overall, these data indicate that exposure to a sublethal CSE dose induces the transcription of cytokines and chemokines, suggesting an active role for bronchial epithelial cells in recruiting neutrophils and macrophages.

## Discussion

COPD is responsible for three million deaths worldwide each year, is caused mostly by cigarette smoking, and has no resolutive therapeutics (1, 2). Excessive airway inflammation and remodeling remain even after several years of smoking cessation, suggesting autoimmunity as a significant driver of the ongoing processes (3, 4). In the last decade, mislocalized mtDNA has been shown to promote inflammatory signaling (11–14). In this study, we wanted to determine whether CS exposure triggers mtDNA release, and whether it is detected in COPD. Measuring extracellular mtDNA and understanding its role may identify novel therapeutic targets for smokers and COPD patients.

Our data revealed elevated levels of cf-mtDNA in the plasma of former smokers with COPD **(Figure 1A-B)** and in the serum of mice with CS-induced enlarged airspace **(Figure 1C-F; supplemental discussion_1)**. To understand the mechanism of its release, we exposed BEAS-2B cells to a sublethal dose of (10%) CSE **(Figure 2A-B)**, and observed decreased mitochondrial membrane potential, increased oxidative stress **(Figure 4A-G)**, and altered mitochondrial dynamics **(Figure 5A-M)**. These changes occurred concomitantly with replicative senescence, as demonstrated by the expression of senescent markers and cell cycle inhibition **(Figure 2C, 4H-L)**. Noteworthy, we described two ways of mtDNA extrusion, by EVs and cell debris **(Figure 3B-E)**. EVs showed a relative increase in mtDNA content over nDNA with increasing CSE doses, while cell debris showed a relative decrease **(Figure 3D, G)**, suggesting a different paradigm of release and signaling.

Necroptosis has been shown as a driving mechanism of cell death caused by exposure to high doses of (16-20%) CSE, and in the lung of COPD patients (7, 21). The upregulation of RIPK3 in COPD lungs and RIPK1 and RIPK3 in CS-exposed murine lungs **(Figure E2)** suggest that necroptosis is a possible mechanism of mtDNA release. However, RIPK1 and RIPK3 were not upregulated in cells exposed to a sublethal (10%) dose of CSE **(Figure E3)**. This is consistent with a recent study that used the same cell line, and in which necroptosis was not driven by upregulation of RIPK3, but by phosphorylation of MLKL (21). Furthermore, in our study, FBS was not included in the media to avoid EV contamination, and its absence may have influenced cell death signaling, including necroptosis.

Our data showing the association of cf-DNA release with CS exposure and COPD matches with other reports of human cohorts. In a small single-center study, higher levels of total cf-DNA (measured by spectrophotometry) were detected in the plasma of patients with COPD exacerbations admitted to the hospital compared to COPD patients without exacerbations and healthy controls, and were associated with an increased risk of 5-year mortality (37). Recently, in a subcohort from the Subpopulation and Intermediate Outcome Measures in COPD Study (SPIROMICS), elevated plasma cf-mtDNA levels were observed in patients with mild and moderate COPD compared to smokers without airflow obstruction (38). Similarly, urine cf-mtDNA levels were associated with increased respiratory symptoms among smokers, and correlated with worse spirometry and chest computed tomography scans in males with emphysema, and with worse respiratory symptoms in females (39).

In our *in vitro* model **(Figure 7)**, CSE exposure induced nDNA release predominantly by cell debris **(Figure 3C, F)**, supporting the idea that it is released by dead cells, as observed in neutrophils (40). On the contrary, mtDNA was released by cell debris and EVs **(Figure 3B, E)**, trended with decreased mitochondrial membrane, increased superoxide, and protein oxidation **(Figure 4)**. In cells exposed to a sublethal dose of CSE, the increased mtDNA in the EVs did not influence the intracellular mtDNA content and TFAM levels **(Figure E3E-H)**, suggesting that the amount of cf-mtDNA released is insufficient to cause detectable depletion among the hundreds of mtDNA copies per cell. Furthermore, at the same sublethal dose of (10%) CSE, 35% of these cells were senescent **(Figure 4I)**. Senescence has been shown to increase the mitochondrial mass (41, 42) and would be expected to offset cellular decreases of mtDNA content.

**Figure 7.**
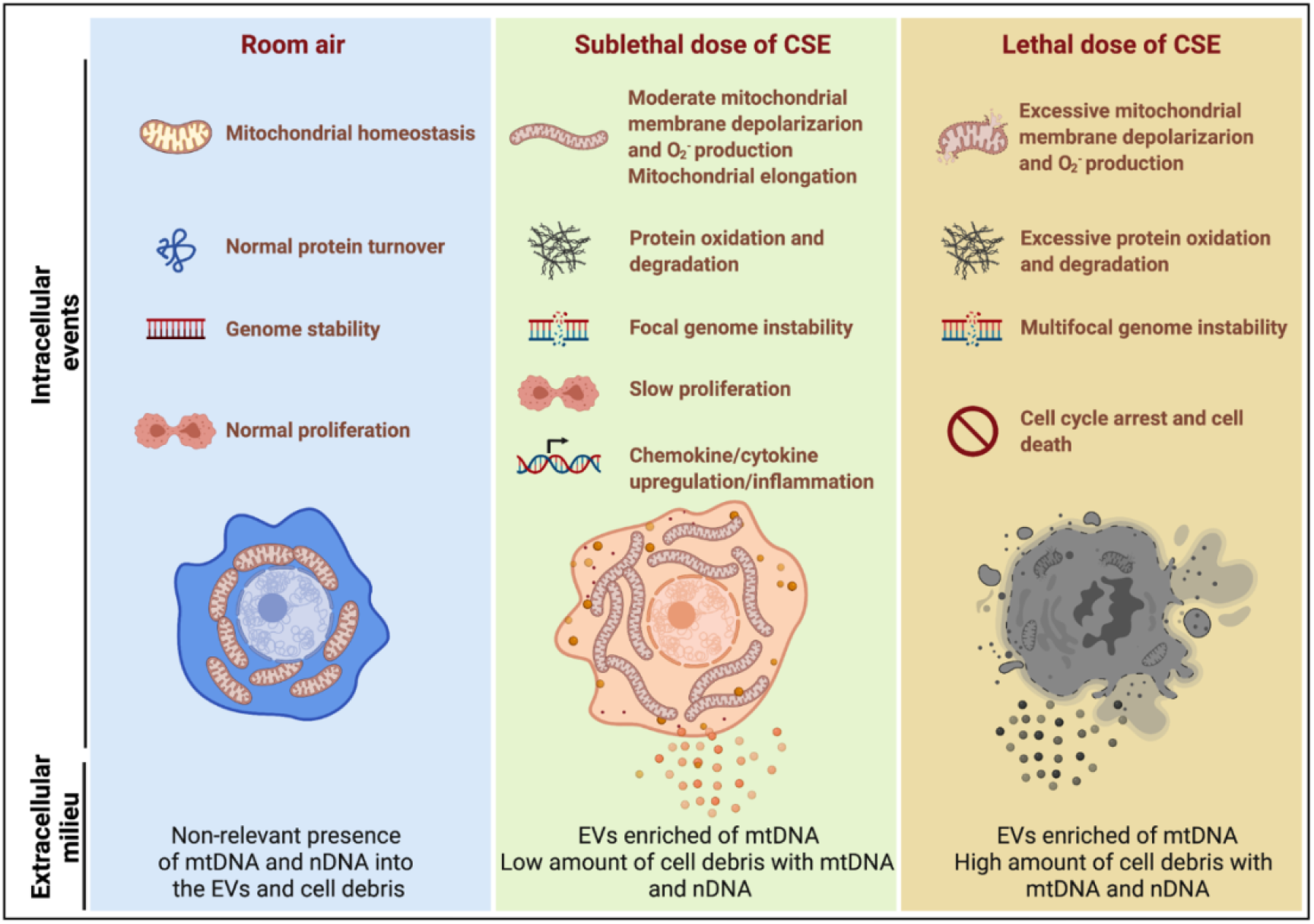
Model summarizing the effects of sublethal (10%) or lethal (20%) doses of CSE with extracellular mtDNA and nDNA release. A sublethal dose (10%) of CSE promoted moderate mitochondrial membrane depolarization, increased superoxide production and oxidative stress, slowed cell proliferation and triggered mtDNA release by extracellular vesicles. This condition is exacerbated in cells exposed to lethal dose (20%) of CSE, in which mtDNA and nDNA are mainly released by the high amount of cell debris.

Interestingly, BEAS-2B cells have been shown to increase exosome release when exposed to CSE. This process is driven by the thiol-reactive properties of CSE, especially acrolein, which may deplete the free thiol group of glutathione (43). Oxidative stress may also alter mitochondrial dynamics, which is one of the mechanisms believed to exacerbate mtDNA packaging into EVs (27). In this regard, we found decreased MFN1, MFN2, and L-OPA in BEAS-2B exposed to a sublethal dose of CSE and in emphysematous human lung tissues **(Figure 5I-J, L-P, supplemental discussion_2)**. Furthermore, a sublethal dose of CSE induced mitochondrial elongation **(Figure 5A-H)**, that has been observed during autophagy as a compensatory mechanism to produce ATP and sustain viability under stressful conditions (44). This process could precede mitochondrial fission (45) and/or may lead to damaged mitochondria (including mtDNA) at the cell periphery (44) to then be extruded in EVs. Recently, it has been shown that a similar mechanism driven by impaired autophagy, leads dysfunctional mitochondria to be extruded from cardiomyocytes, and uptaken and degraded by macrophages (46). On the other hand, because long range PCR suggested the presence of the whole mtDNA in human plasma and EVs **(Figure E1A-C, E)**, it is also plausible to hypothesize an increased horizontal transfer of mtDNA or whole mitochondria between cells to compensate for the bioenergetic defects induced by CS. The function and destination of CSE-induced EVs have not been established.

Two features that we detected, elevated IL-1β and nDNA damage **(Figure 6D, and 2D-F)**, may provide additional insights into the mtDNA mislocalization. It has been shown that IL-1 β can trigger mtDNA release into the cytosol, which stimulates interferon production in the absence of cell death (47). Alternatively, mtDNA release may serve as a genotoxic sentinel of nDNA damage, aiding to preserve the nuclear genome through adaptive gene expression (48) **(supplemental discussion_3)**.

As shown in our model **(Figure 7)**, a (10%) sublethal dose of CSE induced mitochondrial dysfunction, moderate oxidative stress, mtDNA release, and upregulated DSRs (cGAS and NLRP3), DNA responsive genes (DNase III), and proinflammatory cytokines characteristic of COPD inflammation. Although not proven in this case, this suggests that mislocalized mtDNA in cells exposed to a sublethal dose of CSE could contribute to activation of an immune response by cGAS (IL-6), and/or NLRP3 (IL-1β/IL-18), and/or TLR-9 (IL-1β/IL-6/IL-8) signaling (11), and may act as an autocrine and paracrine signal that precede cell death.

The presence of mislocalized DNA could also trigger a compensatory expression of DNases to avoid the immune response. A recent report showed that DNase I treatment could diminish the pathogenic effects of acute CS exposure in mouse lungs (49). This result supports the hypothesis that the extracellular release of DNA following CS exposure contributes to disease etiology. Notably, cf-mtDNA could be the primary driver of the pathophysiological role of cf-DNA induced by CS exposure because of its high copy number (relative to nDNA) **(Figure 1A-B, E-F and 3B-C, E-F)**.

While the present study has demonstrated substantial support for our hypothesis, it has some limitations: i) while our *in vivo* data in emphysematous mice and human former smokers with COPD confirmed high levels of cf-mtDNA in the serum and plasma, our findings need to be validated in a large cohort of patients and intersected in a longitudinal study with clinical parameters to understand the potential impact on the disease. It will also be useful to investigate the presence of cf-DNA in EVs isolated from the plasma and BAL of smokers, COPD patients, and healthy control subjects; ii) CSE is a surrogate for smoke exposure in cultured cells but does not identically recapitulate CS exposure. Patient-derived cells from bronchial brushings may validate our observations; iii) while we used a representative immortalized cell line (BEAS-2B), the airway and lungs host multiple cell types (AECI, AECII, endothelial, smooth muscle, and immune cells) that could be involved in DNA release or its clearance. Future experiments should investigate the role of each cell type; iv) whether extruded mtDNA acts as an inflammatory signal by binding DSRs, or whether it is involved in restoring mitochondrial bioenergetics or is simply a marker for mitochondrial dysfunction in the lung are two unresolved questions. Forthcoming studies should answer these questions by using several knockout cell and animal models.

In conclusion, we demonstrated that CS exposure triggers extracellular mtDNA release in BEAS-2B cells and in a mouse model of emphysema, and we showed that cf-mtDNA levels are elevated in former smokers with COPD. Understanding the mechanism of mtDNA extrusion, and its extracellular role may identify novel therapeutic targets for smokers and COPD patients.

**Supplemental discussions are reported in the online data supplement**

## Acknowledgments

The authors thank Kevin Redding (Heart, Lung, and Blood Vascular Medicine Institute, University of Pittsburgh) for technical assistance and feedback and Meghan Bowler for editorial support.

**Author disclosures** are available with the text of this article at www.atsjournals.org.

## Footnotes

### Author contributions

LG designed and performed the experiments with the help of HH and SAW. ADG isolated serum and lungs from mice and quantified mean chord length, MPV acquired images, and analyzed the immunostained samples. EDeV analyzed flow cytometry data, TB analyzed the EV distributions. YZ and FS provided serum and human lung samples. PS, and SS contributed with feedback and resources to develop the project. LG and BAK conceived and developed the project. LG wrote the manuscript with the input and assistance of all other authors.

