## Supplemental files for "Extracellular release of mitochondrial DNA is triggered by cigarette smoke and is detected in COPD"

#### Supplemental materials and methods

##### Mean chord length (MCL) measurement

The mean alveolar chord length was measured on paraffin-embedded left mouse lungs. Sections were stained with modified Gill's stain, and ten randomly selected 200X fields per slide were analyzed using Scion Image software (version 4.0.2, Scion Corp., Frederick, MD). Airway and vascular structures were masked from the analysis and the images were manually thresholded (1).

##### Cell culture conditions and CSE exposure

BEAS-2B cells (ATCC CRL-9609, Manassas, VA) were maintained in DMEM/F12 (Gibco, Cat#11320-033) supplemented with 5% FBS (Gibco, Cat#10437-028), 100 U/ml penicillin, and 100 μg/ml streptomycin (HyClone, Logan, UT, USA, Cat#SC30010). Trypsin (Thermo Fisher Scientific, Cat#25200056) and a hemacytometer (Hausser Scientific, Horsham, PA, USA, Cat#3200) were used to detach and count cells, respectively. For the experiments, $1x10^5$ cells were plated in a 6-well plate (Genesee Scientific, San Diego, CA, Cat#25105) and cultured overnight, then washed with PBS (Gibco, Cat#10010023) and incubated for 24 hours (hrs) in 2 ml of fresh medium containing cigarette smoke extract (CSE) at the concentrations indicated in the figures. All the experiments in which cells were incubated with CSE (including the unexposed cells, control) were performed in a growth medium without FBS and antibiotics.

Living cells were washed with PBS, detached with 0.025% trypsin for two minutes (mins) at 37˚C, and collected by centrifugation at 1,000 x *g* for five mins. The supernatant was removed and the cell pellet was resuspended in cold PBS, transferred into a new tube, centrifuged at 1,000 x *g* for five mins at 4˚C, and frozen at -80˚C.

**Extracellular vesicle isolation and characterization**

BEAS-2B cells ($1x10^5$ /well) were seeded in 6-well plates and treated the following day with CSE for 24 hrs. One ml of growth medium was collected, centrifuged at 2,000 x *g* for 30 mins to remove the cell debris, then frozen at -80˚C. The supernatant was used to isolate extracellular vesicles (EVs) by the Total Exosome Isolation kit (Invitrogen, Life Technologies, Carlsbad, CA, cat#4478359), as described by the manufacturer. The size and concentration of nanoparticles (*i.e.* nanoparticle tracking analysis) were measured on fresh isolated EVs using a NanoSight NS300 instrument (NanoSight, Malvern Instruments, Malvern, UK). Samples were diluted in PBS to a final volume of 500 µl, then injected into the NS300 unit. Shutter and gain were consistent for the experimental series. A 60-second video was recorded three times for each sample, and then analyzed with the Nanoparticle Tracking Analysis (NTA) 2.0 Analytical software as described elsewhere (2). Size and concentration reports were obtained for each data set.

**Cell free-DNA extraction and quantification**

Cell-free DNA (cf-DNA) was extracted by ethanol precipitation as described previously with minor modifications (3). Briefly, 150 µl of human plasma, or 100 µl of mouse serum, cell debris, EVs, and cell pellets, were thawed and resuspended in proteinase K buffer (100 mM Tris-HCl pH 8.5, 5 mM EDTA, 0.2% SDS, 200 mM NaCl), supplemented with 0.01% 2-mercaptoethanol and 0.6 mg/ml proteinase K (Fisher Scientific, cat# 97062-238) in a final volume of 500 µl, and digested overnight at 55°C. After the addition of NaCl (to the final concentration of 1.27 M) and centrifugation at 28,000 x *g* for 15 mins at 4°C to remove the protein pellet, 800 µl of

100% ethanol and 1 µl GlycoBlue co-precipitant (Thermo Fisher Scientific, cat#AM9516) were added to the supernatant, and nucleic acids were precipitated by centrifugation at 28,000 x *g* for 15 mins at 4°C. The pellet was washed with 70% ethanol, air-dried, and resuspended in DNase-free water. DNA was treated with 0.4 mg/ml RNase A (Sigma-Aldrich, cat#12091021) for 2 hrs at 37°C to degrade RNA, then stored at -20°C.

Cell free-mitochondrial DNA (cf-mtDNA) and cell free-nuclear DNA (cf-nDNA) were quantified by duplex qPCR using TaqMan chemistry. Primer/probe ratios of 1:1 nmole (i.e. primer-limited, PL) for mtDNA assays and 3:1 nmoles for nDNA assays were used, designed on the mitochondrial NADH-ubiquinone oxidoreductase core subunit 1 (ND1PL) and nuclear-encoded ß2-microglobulin (B2M) for human samples (4), and a ND1PL and nuclear-encoded histone deacetylase 1 (HDAC1) probe set for the mouse serum. Primer set and probe sequences are described in **Table E2**. A final volume reaction of 10 µL containing 4 ng of template DNA and 2X Luna Universal qPCR Master Mix (New England Biolabs, Ipswich, ME, USA, cat#M3004) or 2X TaqMan Fast Advanced Master Mix (Thermo Fisher Scientific, cat#4444965) were used. Thermocycling conditions (for both, ND1PL/B2M and ND1PL/HDAC1) were performed in a 96-well StepOnePlus™ Real-Time PCR System (Applied Biosystems, Thermo Fisher Scientific, cat#4376600) as described: an initial denaturation step at 95°C for 20 seconds (s), followed by 40 cycles each consisting of a denaturation step at 95°C for 1 s, an annealing step at 63°C for 20 s, and an elongation step at 60°C for 20 s. Serial dilutions of DNA were used to ensure assay linearity, and to calculate the relative abundance of individual samples. Absolute copies/µl of DNA were determined by interpolation of the standard curve analyzed by singleplex Digital PCR using the QuantStudio 3D Digital PCR System (Applied Biosystem, Thermo Fisher

Scientific) for each primer set. Digital PCR reactions were performed in duplicate (two chips per primer).

**Cytotoxicity assay and β-Galactosidase staining**

Cytotoxicity screening was conducted by cell density determination using a sulforhodamine B (SRB) assay to measure total cellular protein content as described previously (5, 6) with some modifications. Briefly, $7x10^4$ cells were plated in a 24-well plate overnight. Then, cells were washed with PBS and incubated for 24 hrs with CSE. The cell monolayer was fixed with 10% cold trichloroacetic acid (Sigma-Aldrich, St. Louis, MO, USA, cat#T6399) at 4ºC for 1 hour (h), washed with PBS and dried at 37ºC for 90 mins in a heater (Isotemp, Thermo Fisher Scientific). Cells were stained with 0.4% SRB solution (Sigma-Aldrich, cat#S1402) for 20 mins at room temperature (RT) on a shaker. Non-specific SRB staining was removed by five washes with 1% acetic acid, and the cell monolayer was dried at 37ºC for 90 mins in the heater. The protein-bound dye was dissolved by adding 10 mM Tris (pH 10.5) to the well followed by gentle shaking for 10 mins. Two-hundred µl of the dissolved solution was transferred to a 96-well plate and the absorbance was read in triplicate at 510 nm at 25°C using a Synergy H1 (BioTek) microplate reader.

β-galactosidase (β-gal) staining was performed using the Senescence β-Galactosidase Staining Kit (Cell Signaling Technology, MA, USA, cat#9860), following the manufacturer's instructions. Briefly, $1x10^5$ cells/well were seeded in a 6-well plate overnight. Cells were treated with CSE for 24 hrs, followed by fixation for 15 mins, rinsed with PBS, and incubated with the β-gal staining solution at 37°C overnight. After rinsing with PBS, the crystals were solubilized in 50% DMSO and images were taken by an Axiovert 40 CFL microscope (Carl Zeiss, White Plains, NY,

USA) using an AxioCamMRC5 camera and AxioVision LE 4.8 software. β-gal positive cells were quantified using the cell counter plugin by ImageJ (7). The ratio of β-gal-positive cells to the total number of cells per single image was expressed as a percentage. Images were adjusted for brightness and contrast, and resized and annotated for presentation without any further manipulation.

**Mitochondrial membrane potential and superoxide production measurements**

Mitochondrial membrane potential and superoxide production were measured by Tetramethylrhodamine ethyl ester (TMRE; Abcam, Cambridge, UK, cat#ab113852, Abcam) and MitoSOX Red (Thermo Fisher Scientific, cat#M3600) probes, respectively. A Sinergy H1 microplate reader (BioTek) was used to measure fluorescence, set at 549 nm $\lambda_{ex}$ and 575 nm $\lambda_{em}$ for TMRE, and 510 nm $\lambda_{ex}$ and 580 nm $\lambda_{em}$ for MitoSOX. Briefly, $7x10^5$ cells were seeded in a 10 cm dish. After 24 hrs, the cells were washed with PBS and incubated with increasing concentrations of CSE for 3 or 6 hrs, as indicated in the figure legend. Then, the cells were washed with PBS, detached with trypsin, and centrifuged at 1,000 x *g* for 5 mins.

To measure the mitochondrial membrane potential, cells were resuspended in fresh media containing CSE supplemented with 250 nM TMRE, and incubated for 25 mins at 37°C. Next, cells were centrifuged at 1,000 x *g* for 5 mins and resuspended in 1 ml of 0.02% bovine serum albumin (BSA; Sigma-Aldrich, cat#A9418) in 1X PBS. Fluorescence was measured using $1.4x10^4$ cells/well in a 96-well black plate (Greiner Bio-One, Monroe, NC, USA, cat#655076). Cells incubated without TMRE or with 20 µM carbonyl cyanide p-trifluoro-methoxyphenyl hydrazone (FCCP) were used as negative and positive controls, respectively.

To measure superoxide production, the fluorescence variation (ΔRFU) during a time course of 10 mins was measured in $1.4x10^5$ cells/well using a 96-well black plate (Greiner Bio-One, cat#655076) in cells previously resuspended in HBSS (Gibco, 14175) and probed with 5 µM MitoSOX. Cells pretreated with 100 µM MitoTEMPO (Enzo Life Sciences, Farmingdale, NY, USA, cat#ALX-430-150-M005), or incubated with 50 µM Antimycin A (Sigma-Aldrich, A8674) were used as negative and positive controls, respectively.

**Cell cycle analysis**

Cell cycle arrest was quantified by detecting the incorporation of 5-ethynyl-2'-deoxyuridine (EdU) during *de novo* DNA synthesis using the Click-iT EdU Alexa Fluor 488 Flow Cytometry assay Kit (Thermo Fisher Scientific, cat#C10425). Briefly, cells were co-incubated with 2 µM EdU the last 2 hrs of CSE exposure, then they were collected and processed following the manufacturer's instructions, and analyzed by an LSR II flow cytometer (BD Biosciences, San Jose, CA, USA). EdU-free cells were used as a negative control.

**Immunocytochemistry**

$1x10^5$ cells were plated in a 6-well plate and cultured overnight on a 35 mm glass coverslip (MatTEK Corporation, Ashland, MA, USA, cat#P35GCOL-1.5-10-C). Then they were incubated with CSE for 24 hrs, washed with PBS, fixed with 4% paraformaldehyde (Electron Microscopy Sciences, Hatfield, PA, USA, cat#15719-S) for 20 mins at RT, washed again with 0.3 M glycine-PBS (3 times, 5 mins each), permeabilized with 0.3 M glycine-0.1% Triton-X100 for 15 mins at RT, and blocked for 30 mins with 0.3 M Glycine 0.05% Triton-X100, 0.5% BSA (blocking solution).

Cells were stained with primary antibodies diluted in blocking solution overnight at 4°C as follows: rabbit anti-53BP1, (1:250, Abcam, cat#175933), mouse anti-α-tubulin (1:1000, Sigma Aldrich, cat#T6074), and anti-TOMM20 (1:250, Thermo Fisher Scientific, cat#PA5-52843). After rinsing with PBS (3 times, 5 mins each), cells were probed for 1 h, at RT temperature with the following secondary antibodies: Alexa Fluor 647 goat anti-rabbit (1:500, Thermo Fisher Scientific, cat#A21245) and Alexa Fluor 488 goat anti-mouse (1:500, Thermo Fisher Scientific, cat#A2817) diluted in blocking solution, followed by a wash in PBS (3 times, for 5 mins). Nuclei were stained using 2 μg/ml Hoechst for 20 mins (bis-benzimide H33258, Sigma Aldrich, cat#B2883) at RT. Coverslips were mounted by using Gelvatol medium on a microscope slide (Thermo Fisher Scientific, cat#1255015).

**Spinning disk confocal microscopy and image analysis**

All images were acquired using a spinning-disk Marianas system based on a Zeiss Axio Observer Z1 inverted fluorescence microscope equipped with a 63X Plan Apo PH NA 1.4 oil immersion objective, Evolve EM-CCD camera, piezo stage controller, spherical aberration correction module and lasers (405, 488, 515, 561 and 640 nm), all of which were controlled by SlideBook6 software (Intelligent Imaging Innovation, Denver, CO). A z-stack of fifteen 2D-confocal images were acquired in 400 nm steps. Image acquisition settings were identical for all variants in each experiment, using the 405 nm laser (for Hoechst), 488 nm laser (for Tubulin), and 640 nm laser (for 53BP1 and TOM20).

To count the 53BP1 foci, 3D images were deconvolved using the constrained iterative algorithm of SlideBook6. A segment mask was then generated to select 53BP1 puncta detected through the 640 nm channel (53BP1 mask), and objects smaller than 5 voxels were eliminated. Another segment mask was generated with the minimum threshold in the 405 nm channel to include total Hoechst fluorescence (Hoechst mask), and objects smaller than 100 voxels were eliminated. For both masks, identical threshold parameters were used for all experimental variables. The number of objects in 53BP1 mask was divided by the number of objects in the Hoechst mask and reported as the number of foci/nucleus. The volume of each object of the 53BP1 mask was also calculated. Six-eight images per condition containing multiple cells were analyzed for each independent experiment.

Mitochondrial morphology analysis was carried out using maximum projection of 3D images, after background subtraction. A segment mask was created for the 640 nm channel to select the TOMM20 signal (TOMM20 mask), and objects smaller than 5 voxels were eliminated from the mask. Another segment mask was generated with the minimum threshold in the 405 nm channel to include total Hoechst fluorescence (Hoechst mask), and objects smaller than 100 voxels were eliminated. Incomplete or intertwined cells were manually removed from both masks. The number of objects (TOMM20-positive mitochondria), the total area, and the total perimeter of the TOMM20 mask were divided by the number of objects of the Hoechst mask to calculate the number of mitochondria, mitochondrial area, and perimeter per cell. For the mitochondrial morphological analysis, the area, perimeter, aspect ratio (major axis/minor axis of the ellipse equivalent to the object), and form factor ($perimeter^2/4\pi \cdot area$; inverse of sphericity) were calculated for each object of the TOMM20 mask as previously described (8). Six-eight images per condition containing multiple cells were analyzed for each independent experiment.

**Long-range PCR**

Long-range PCR was performed as previously described (9) with minor modifications. Briefly, DNA extracted from human plasma (2.5 ng) or EVs (3 ng) was amplified using 0.5 µM primers and LongAmp Hot Start Taq 2X Master Mix (New England BioLabs, cat#M0533S) in a final volume of 50 µl by a ProFlex PCR System Thermocycler (Applied Biosystems). Thermocycling conditions were set as follows: polymerase activation (94°C for 2 mins), then 30 cycles, each consisting of denaturation (94°C for 15 seconds), annealing and elongation (62°C for 12 mins), with a final extension stage (72°C for 10 mins). PCR products were run on a 0.7% agarose gel. Primers used for this assay were previously described (9, 10) and listed in **Table E3**.

**RNA extraction and RT-qPCR**

RNA was extracted from $1.5 \times 10^5$ cells using the RNeasy Mini Kit (Qiagen, Valencia, CA, USA, cat#74104) and treated with DNase I to remove DNA contamination using the DNA-free Kit DNase Treatment and Removal Reagents (Life Technologies, cat#AM1906) following the manufacturer's instructions. RNA was quantified by NanoDrop (Thermo Fisher Scientific, NanoDrop One$^{C}$) and cDNA was synthesized using 1 µg of RNA by High-Capacity cDNA Reverse Transcription Kit (Applied Biosystems, cat#4368813) in a ProFlex PCR System Thermocycler (Applied Biosystems). Real-time qPCR was carried out using 2 µl of cDNA as template (coming from the RT-qPCR diluted 1:10 or 1:100), 0.4 µM primers, and Power Sybr Green PCR Master Mix (Applied Biosystems, cat#4367659) in a final volume of 10 µl, using a 96-well StepOnePlus™ or a 384-well QuantStudio5 Real-Time PCR System (Applied Biosystems). Primer sets were designed using the Primer Blast software (https://www.ncbi.nlm.nih.gov/tools/primer-blast) and are reported in **Table**

**E4**, or were previously described (11). Quantitative PCR was run with an initial UDG activation at 50°C for 2 mins, denaturation at 95°C for 10 mins, followed by 40 cycles of denaturation (95°C, 15 seconds) and annealing and synthesis (59°C, 1 min). Fold change was calculated by the $2^{-\Delta\Delta Ct}$ method using β-Glucuronidase (GUSB) as a housekeeping gene.

**Protein extraction, SDS-PAGE, and Western blotting**

Snap-frozen human and mouse lung tissues were mechanically homogenized by a pestle in a cold mortar submerged in liquid nitrogen vapor. Twenty-five mg of powder were dissolved in modified RIPA buffer ((150 mM NaCl, 50 mM Tris-HCl, pH 7.4, 0.25% sodium deoxycholate, 1 mM EDTA, 1% NP-40, 1% complete protease inhibitor (Thermo Fisher Scientific, cat#36798), and 1% phosphatase inhibitor (VWR, G-Biosciences, St. Louis, MO, USA, cat#95029-242)) on a rotator at 4°C for 30 mins. Cell pellets were also lysed in RIPA buffer on ice for 30 mins. Lysates were centrifuged at 20,000 x *g* for 10 mins at 4°C, and the supernatant was transferred to pre-chilled tubes for protein quantification using the Micro BCA Protein Assay (Life Technologies, cat#23235). Ten or fifteen µg of protein were separated on a 4-12% bis-tris polyacrylamide gel (Thermo Fisher Scientific, cat#NW04122), transferred to a nitrocellulose membrane using an iBlot gel transfer device (Invitrogen, Life Technologies), blocked with 0.25X Intercept Blocking buffer (Li-COR Biosciences, Lincoln, NE, USA, cat#927-40000), and immunoblotted overnight at 4°C using antibodies against the following proteins: GAPDH (1:2000, Cell Signaling Technology, cat#2118), MFN1 (1:2000, Cell Signaling Technology, cat#14739), MFN2 (1:2000, Cell Signaling, cat#9482), MLKL (1:1000, Cell Signaling Technology, cat#14993), OPA1 (1:1000, Cell Signaling Technology, cat#80471), p21

(1:1000, Cell Signaling Technology, cat#2947), p62 (1:1000, Sigma Aldrich, cat#PA0067), RIP1 (1:4000, Cell Signaling Technology, cat#3493), RIP3 (1:2000, Abcam, cat#56164), TFAM (1:2500, PhosphoSolutions, Aurora, CO, USA, cat#1999), Ubiquitin (1:1000, Cell Signaling Technology, cat#3933), and β-actin (1:3000, Santa Cruz Biotechnology, Dallas, TX, USA, cat#sc47778) diluted in 0.25X Intercept Blocking buffer with 0.02% Tween 20. The membrane was washed (3 times, 10 mins each) with TBST (TBS, 0.001% Tween 20) and incubated for 1 h at RT with the following secondary antibodies: goat anti-mouse Alexa Fluor 680 (1:10000, Thermo Fisher Scientific, cat#A21058) and goat anti-rabbit Alexa Fluor 800 (1:10000, Thermo Fisher Scientific, cat#A32735). The membrane was washed again with TBST (3 times, 10 mins each), and fluorescence was detected on a ChemiDOC MP Imaging System (Bio-Rad Laboratories, Hercules, CA, USA). After detection, the membrane was incubated with stripping solution (Li-COR Biosciences, cat#92840030) for 25 mins at RT, washed with 1X TBS (2 times, 10 mins each), and re-probed overnight.

Carbonylated proteins were detected using the Oxyblot Protein Oxidation Detection Kit (Millipore, Sigma-Aldrich, cat#S7150) following the manufacturer's instructions. Briefly, proteins were extracted as described above, except for the addition of 5% β-mercaptoethanol in the RIPA buffer to prevent oxidation. Equal amounts of protein (8 μg) were mixed with 5% SDS, then the carbonyl groups in the protein side chains were derived by incubating with 2,4-dinitrophenylhydrazine (DNPH) for 15 mins at 30°C to generate 2,4-dinitrophenylhydrazone (DNP hydrazone). After blocking the reaction with neutralization solution for 5 mins, protein samples were run on a 4-12% SDS-polyacrylamide gradient gel followed by western blotting that was performed as described above, except that the blocking agent was 3% BSA in PBS 1X, 0.1% Tween

20 (PBST). The membrane was incubated overnight with an antibody against DNP-hydrazone (included in the kit, diluted 1:400 with 1% BSA-PBST). After washing with PBST (3 times, 10 mins each), the membrane was incubated for 1 h with the secondary antibody (included in the kit, diluted 1:1000 in 3% BSA PBST), and washed again with PBST (3 times, 10 mins each). The immunoreactivity was detected using the chemiluminescent substrate Pierce ECL Plus (Thermo Fisher Scientific, cat#32132) on a ChemiDOC MP Imaging System (Bio-rad Laboratories). After detection, the membrane was stripped as described above and re-probed overnight with an anti-GAPDH antibody (rabbit, 1:2000, Cell Signaling Technologies, Cat#2118S) in 1% BSA-PBST. After washing with PBST (3 times, 10 mins each), the membrane was incubated with the secondary antibody goat anti-rabbit (1:12500, Santa Cruz Biotechnology cat#sc-2004) diluted in 1% BSA PBST at RT for 1 h, followed by washes with PBST (3 times, 10 mins each). The GAPDH signal (previously described) was detected and used to normalize the total carbonylated proteins.

**Illustration**

The schematic illustration in Figure 7 was created using BioRender.com.

#### Supplemental tables

**Table E1. Clinical and demographic characteristics of the human plasma study cohort**

| | Former Smoker | Former Smoker COPD | P-value |
|---|---|---|---|
| **N** | 14 | 20 | |
| **Age (years)** | 63.5 ± 6.5 | 59.9 ± 5.1 | 0.006 |
| **Gender (% Female)** | 57.10% | 75.00% | 0.2874 |
| **Packs per year** | 48.6 ± 23.8 | 49.3 ± 36.1 | 0.9556 |
| **$FEV_1$% Predicted** | 99.5% ± 10.2% | 23.0% ± 3.4% | <0.0001 |
| **$FEV_1$/FVC** | 0.793 ± 0.03 | 0.318 ± 0.06 | <0.0001 |
| **DLCO% Predicted** | 91.6% ± 12.7% | 29.7% ± 9.0% | <0.0001 |
| **LAA%** | 0.004 ± 0.002 | 0.277 ± 0.121 | <0.0001 |
| **Hist 15** | -884.2 ± 23.6 | -964.6 ± 15.4 | <0.0001 |

$FEV_1$, forced expiratory volume in 1 second; FVC, forced vital capacity; DLCO, diffusing capacity of carbon monoxide; LAA%, percentage of low-attenuation area less than −950 Hounsfield Unit on a CT scan; Hist 15, the lower 15th percentile of the Hounsfield Unit value histogram. P-values were determined by an unpaired parametric t-test.

**Table E2. Primers and probes used for mtDNA and nDNA quantification**

| | Primers Probes | 5'-3' sequence |
|---|---|---|
| **human ND1PL** | F | gagcgatggtgagagctaaggt |
| | R | ccctaaaacccgccacatct |
| | Probe | 5HEX/ccatcaccc/ZEN/tctacatcaccgccc/3IABKFQ |
| **human B2M** | F | tctctctccattcttcagtaagtcaact |
| | R | ccagcagagaatggaaagtcaa |
| | Probe | 6-FAM/atgtgtctg/ZEN/ggtttcatccatccgaca/3IABKFQ |
| **mouse ND1PL** | F | gagtgatagggtaggtgcaataa |
| | R | ccatttgcagacgccataaa |
| | Probe | 5HEX/agaaccaat/ZEN/acgccctttaacaacctct/3IABKFG |
| **mouse HDAC1** | F | ccaggtactcgttagtgttctg |
| | R | ctttgaatactttggaccggattt |
| | Probe | 56-FAM/acatcagcc/ZEN/cttccaacatgacca/3IABKFQ |

F and R indicate forward and reverse primers, respectively.

**Table E3. Primer sets used for long-range PCR**

| Primers | | 5’-3’ sequence | Amplicon | Reference |
|---|---|---|---|---|
| **Mito 1** | 16331 F | acatagcacattacagtcaaatcccttctcgtcccc | 3968 bp | Sansone P. et al., 2017 |
| | 3729 R | tgagattgtttgggctactgctcgcagtgc | | |
| **Mito 2** | 3646 F | tactcaatcctctgatcagggtgagcatcaaactc | 5513 bp | |
| | 9458 R | gcttggattaaggcgacagcgatttctaggatagt | | |
| **Mito 3** | 8753 F | tcatttttattgccacaactaacctcctcggactc | 7813 bp | |
| | 16566 R | cgtgatgtcttatttaaggggaacgtgtgggctat | | |
| **Mito 4** | 14841 F | tttcatcatgcggagatgttggatgg | 8842 bp | Furda AM et al., 2014 |
| | 5999 R | tctaagcctccttattcgagccga | | |
| **Mito 5** | 14841 F | tttcatcatgcggagatgttggatgg | 221 bp | |
| | 14620 R | ccccacaaaccccattactaaaccca | | |
| F and R indicate forward and reverse primers, respectively; bp, base pair. | | | | |

**Table E4. Primer sets used for gene expression quantification**

| NM | Gene | Primer | 5’-3’ sequence |
|---|---|---|---|
| NM_138441.3 | cGAS | F | aggcctgcgcattcaaaact |
| | | R | tgtgagagaaggatagccgcc |
| NM_005223 | DNase I | F | ccagacacctatcactacgtgg |
| | | R | ctctcggttgaaggtgtcgttc |
| NM_001375 | DNase II | F | gtcaagggccaccacgttag |
| | | R | gccaaccagccggagtacag |
| NM_033629 | DNase III | F | agagtgtgcagccgagtcac |
| | | R | ccagtggcctccatgtcgaa |
| NM_000181 | GUSB | F | ctgtcaccaagagccagttcct |
| | | R | ggttgaagtccttcaccagcag |
| NM_001562.4 | IL-18 | F | gatagccagcctagaggtatgg |
| | | R | ccttgatgttatcaggaggattca |
| NM_004895.5 | NLRP3 | F | ggactgaagcacctgttgtgca |
| | | R | tcctgagtctcccaaggcattc |
| NM_138554.5 | TLR-4 | F | ccctgaggcatttaggcagcta |
| | | R | accctgaggcatttaggcagcta |
| F and R indicate forward and reverse primers, respectively. | | | |

**<u>Supplemental discussion</u>**

**_1. Cf-DNA detection in mouse models of lung inflammation**

Similar to our results in the mouse model **(Figure 1E-F)**, increased extracellular mtDNA or cf-DNA (measured by fluorescence) were reported in the BAL of mice with acute lung inflammation induced by (1-4 days) short term exposure to CS, even though these models do not mimic emphysema.

In C57BL/6J mice, acute exposure to CS (10 cigarettes) enriched the BAL with mtDNA, as early as 30 minutes after exposure. nDNA was found 10 days later, most likely as a secondary effect of tissue necrosis (12). In the same study, BEAS-2B cells exposed to prolonged low-level oxidative stress induced by incubation with glucose oxidase showed moderate bioenergetic defects, mtDNA extrusion by EVs, and a proinflammatory response characterized by increased IL-1β, IL-6, and IL-8 expression (12). Likewise, in our study, we found increased oxidative stress **(Figure 4A-D)**, mtDNA release by EVs **(Figure 3B**), and upregulated proinflammatory cytokines **(Figure 6C-D)** when we exposed the same cell line to CSE, a model that more accurately mimics CS exposure than glucose oxidase.

Other work, using a different paradigm of acute CS exposure (four cigarettes, three times a day for four consecutive days) to induce mouse lung inflammation, detected double-strand DNA (ds-DNA) by PicoGreen fluorescent dye in the BAL of C57BL/6J mice. Importantly, the lungs of CS-exposed mice, showed upregulated cGAS/STING (stimulator of interferon genes) and type I interferon (IFN I) signaling, together with neutrophil recruitment (13). ds-DNA was also detected in the BAL of other mouse strains (BALB/cByJ, DBA/1J, including C57BL/6J) except for C58/J, acutely exposed (3-5 cigarettes, five days) to CS (14).

**_2. Alteration of mitochondrial dynamics in cells and human lung tissue**

Our results agree with a recent report that showed decreased L-OPA1 protein in several cell lines (including BEAS-2B cells) exposed to acute CSE for 4 hours (15). Furthermore, decreased MFN1 protein was detected in the mitochondrial fraction obtained from lung tissue isolated from smokers without emphysema compared to non-smokers, and in lung homogenates isolated from an area with severe emphysema compared to an area with mild emphysema in the same patient (16).

**_3. Alternative pathways that could induce mtDNA extrusion in cells exposed to CSE**

IL-1β induced mtDNA release into the cytoplasm of type II alveolar cells (AECII) and macrophages, which subsequently activated the innate immune response through the cGAS/STING/interferon regulatory factor 3 (IRF3) cascade (17). Recently, it has also been proposed that mtDNA release into the cytoplasm could enhance nDNA repair responses, acting as a genotoxic stress sentinel for nDNA damage (18). Indeed, we showed that cells treated with a sublethal dose (10%) of CSE had increased IL-1β expression **(Figure 6D)**, nDNA damage and a p16-/p21-dependent cell cycle arrest **(Figure 2D-F, and 4J-L)**. Similarly, other findings have described a p21-dependent and p53-independent cell cycle arrest in AECII from human emphysematous lungs (19).

#### Supplemental figure legends

**Figure E1. Human plasma of former smokers with and without COPD, and EVs released from BEAS-2B cells unexposed and exposed to CSE contain overlapping fragments of the mitochondrial genome**

**(A)** Schematic and **(B-C)** representative images of agarose gel electrophoresis (AGE) of long-range PCR products amplified by five overlapping primer sets spanning the entire mitochondrial genome (Mito1-Mito5). The DNA template was extracted from a random subgroup of former smokers without COPD (n=4; ID: 15292, 33724, 34478, 34663) and former smokers with COPD (n=4; ID15299, 15466, 33229, 15470). **(D)** Distribution of the EVs released into the extracellular milieu by BEAS-2B cells exposed to increasing doses of CSE. **(E)** Representative image of AGE of long-range PCR products (Mito1-Mito5) performed using DNA extracted from EVs isolated from BEAS-2B cells unexposed or exposed to 10% and 20% CSE (n=2 per group). **(D)** Data are reported as mean of three independent experiments (n=3).

CSE, cigarette smoke extract; control, CSE-unexposed cells; EVs, extracellular vesicles; PCR, polymerase chain reaction.

**Figure E2. Necroptosis markers are dysregulated in the airways of COPD patients and in the lungs of a mouse model of emphysema induced by CS exposure**

**(A, D)** Representative western blots showing RIP1 and RIP3 proteins and their relative densitometry **(B, C)** in human airways without signs of lung obstruction (Donor, n=12) or affected by COPD (COPD, n=12), and in **(E-F)** lung tissue of mice exposed to cigarette smoke (n=5) or room air (n=6) for six months. β-actin was used as a loading control. (D) The dotted arrow indicates the cropping of unrelated samples run on the same gel. Data are reported as mean±SEM of several samples (n) indicated as circles. ns, no statistical significance; *p<0.05, or **p<0.01 were determined by an unpaired t-test analysis with Welch's correction.

**Figure E3. Exposure to a sublethal dose of CSE induces superoxide production that is neutralized by MitoTEMPO, but does not increase RIPK1, RIPK3 proteins, and the markers of mitochondrial mass (mtDNA and TFAM content)** (**A)** Bar graph reporting the superoxide anion production determined by MitoSOX as the variation of the relative fluorescence unit (RFU) for 10 mins from BEAS-2B cells previously exposed to CSE for 3 hrs, as reported in Figure 4B. Treatment groups include cells pretreated for 5 hrs with 100 μM MitoTEMPO or incubated with 50 μM Antimycin A, which were used as a negative and positive control, respectively. Representative western blots of **(B)** necroptosis markers (RIPK1, RIPK3, MLKL) and **(G)** TFAM in BEAS-2B cells exposed to increasing doses of CSE. Bar graphs reporting the densitometry of (**C)** RIPK1, **(D)** RIPK3, **(E)** MLKL and **(H)** TFAM proteins, and the **(F)** intracellular mitochondrial DNA content. β-actin was used as a loading control. **(A, C-F, H)** Data are reported as mean±SEM of independent experiments (n≥3), indicated as circles. CSE, cigarette smoke extract; control, CSE-unexposed cells. ns, no statistical significance; *p<0.05, or **p<0.01 were determined by **(A)** unpaired t-test with Welch's correction, **(C-F, H)** one-way ANOVA with multiple comparisons with Bonferroni post hoc test.

### Supplemental figures

**Figure E1**

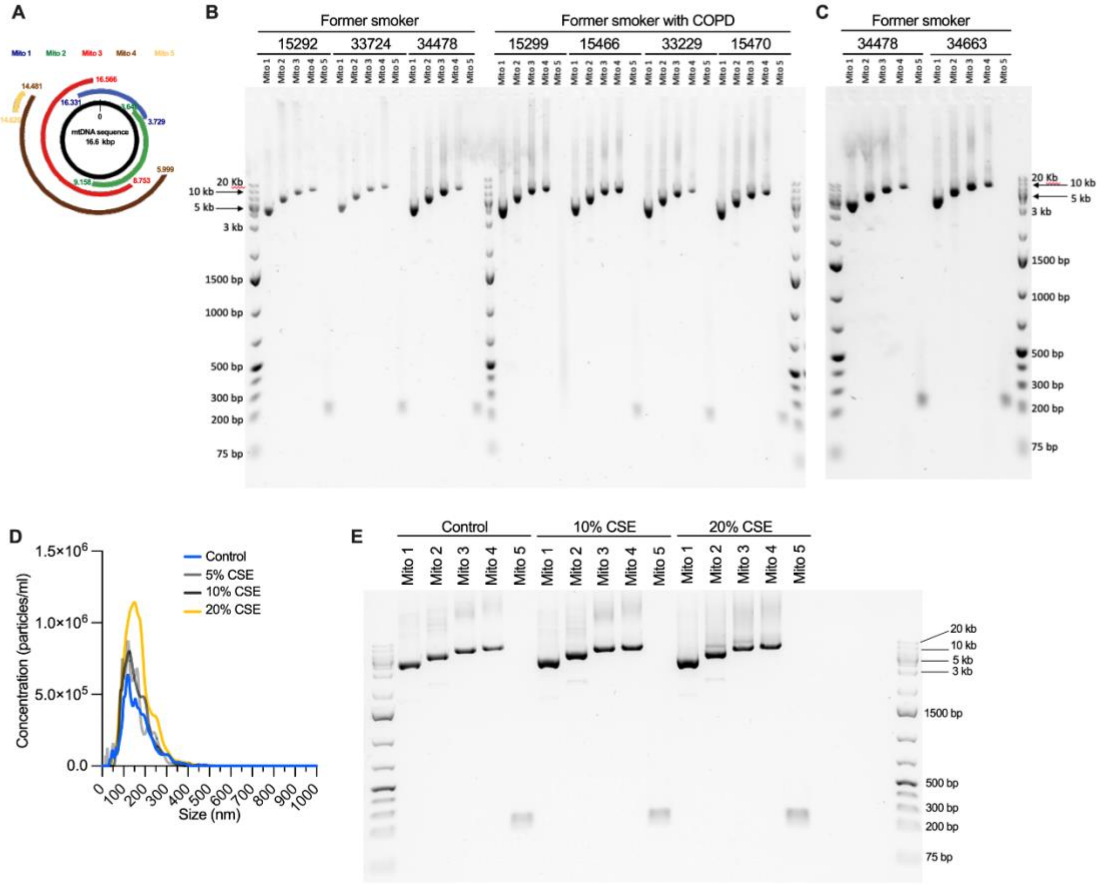

**Figure E2**

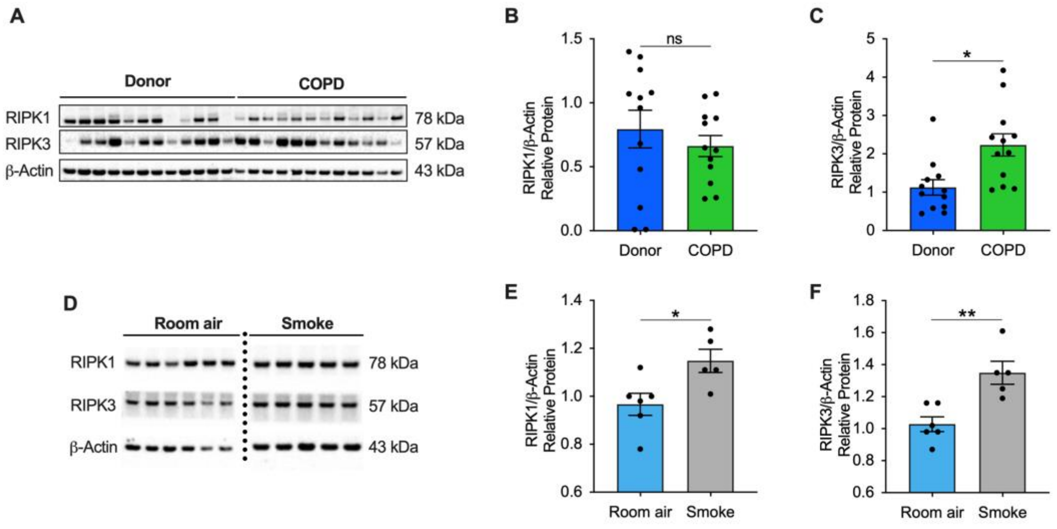
A
Donor
COPD
RIPK1
RIPK3
β-Actin
78 kDa
57 kDa
43 kDa
B
RIPK1/β-Actin Relative Protein
ns
Donor
COPD
C
RIPK3/β-Actin Relative Protein
*
Donor
COPD
D
Room air
Smoke
RIPK1
RIPK3
β-Actin
78 kDa
57 kDa
43 kDa
E
RIPK1/β-Actin Relative Protein
*
Room air
Smoke
F
RIPK3/β-Actin Relative Protein
**
Room air
Smoke

**Figure E3**

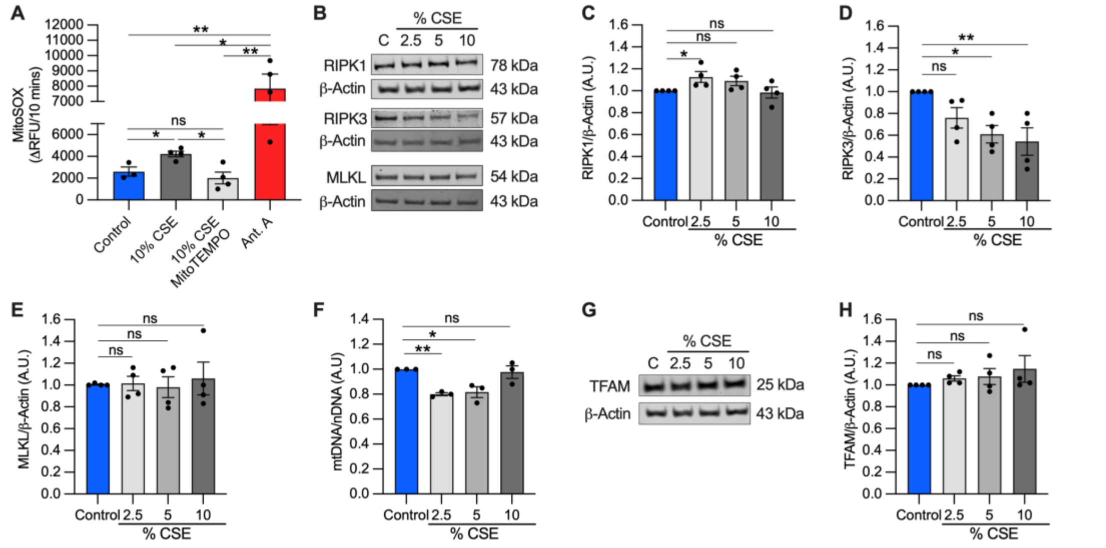
A
MitoSOX (ΔRFU/10 mins)
Control
10% CSE
10% CSE MitoTEMPO
Ant. A
ns
*
**
B
% CSE
C 2.5 5 10
RIPK1 78 kDa
β-Actin 43 kDa
RIPK3 57 kDa
β-Actin 43 kDa
MLKL 54 kDa
β-Actin 43 kDa
C
RIPK1/β-Actin (A.U.)
D
RIPK3/β-Actin (A.U.)
E
MLKL/β-Actin (A.U.)
F
mtDNA/nDNA (A.U)
G
% CSE
C 2.5 5 10
TFAM 25 kDa
β-Actin 43 kDa
H
TFAM/β-Actin (A.U.)
Control 2.5 5 10
% CSE
